# DiffTernary: a Diffusion Model for Molecular Glue Ternary Complex Docking

**DOI:** 10.64898/2026.11.09.765352

**Authors:** Yiyan Liao, Letian Lin, Jintao Zhu, Luhua Lai, Jianfeng Pei

## Abstract

Molecular glues (MGs) modulate protein function by inducing or stabilizing protein interactions. However, accurately modeling MG ternary complexes remains challenging. Here, we present DiffTernary, a diffusion-based docking method for the joint prediction of relative protein arrangement and MG binding pose. We introduce Ternary-PDB, a dataset of 76,702 experimentally determined ternary complexes curated from the Protein Data Bank using structural filters and a release cutoff aligned with AlphaFold 3 (AF3). When trained on this dataset, DiffTernary achieved acceptable protein–protein interaction (PPI) predictions (DockQ > 0.23) for 50.0% of MGBench complexes with a single prediction, compared with 51.1% for AF3. In comparisons with AF3, OpenFold3, ESMFold2, Protenix-v1 and Protenix-v2, DiffTernary ranked second with a best-of-25 PPI prediction success rate of 78.4%, behind ESMFold2 (90.9%) and ahead of AF3 (72.7%). A separate DiffTernary model trained on TernaryDB achieved a mean DockQ of 0.306 on the 94 complexes in the TernaryDB MGD test set with a single prediction, compared with 0.228 for DeepTernary. Confidence-based selection among 40 samples yielded acceptable PPI predictions for 41.5% of complexes, compared with 30.9% for DeepTernary. An E2Former variant increased training and sampling throughput 6.90-fold and 4.37-fold, respectively, under the evaluated configurations. Together, DiffTernary and Ternary-PDB provide an efficient docking approach for MG complex prediction and a valuable structural resource for MG discovery and design.

## Introduction

MGs are typically small, monovalent molecules that induce new protein–protein interactions (PPIs) or stabilize existing ones through ternary complex formation^1–4^. Based on their mechanisms of action, MGs can be classified as MG degraders (MGDs), nondegradative heterodimerizers, or homodimerizers^2,5^. MGDs recruit target proteins to E3 ubiquitin ligases, leading to ubiquitination and proteasomal degradation. Nondegradative heterodimerizers stabilize PPIs without inducing degradation, whereas homodimerizers promote protein self-association^3,6^. Unlike proteolysis-targeting chimeras (PROTACs), which consist of two protein-binding moieties connected by a linker, MGs generally have a compact, monovalent structure. This property makes MGs attractive for targeting traditionally undruggable proteins, including transcription factors and intrinsically disordered proteins, while potentially retaining favorable drug-like properties^1,2^.

Despite increasing interest in MGs, their discovery still relies largely on phenotypic screening, chemical proteomics, and biochemical assays^2,7,8^. Structures of MG ternary complexes reveal how the MG binds and how the two proteins interact, providing a basis for understanding MG activity and guiding molecular design. However, determining MG ternary complex structures by X-ray crystallography or cryo-electron microscopy remains challenging, and relatively few experimental structures are available^3,5,9^. Computational structure prediction can complement experimental studies and support structure-guided optimization and virtual screening of MGs^3,10^. Cofolding methods predict complex structures from protein sequences and molecular representations, whereas docking methods assemble complexes from the structures of their individual components. Cofolding models, including RoseTTAFold All-Atom, AF3, Chai-1, Boltz-1, Boltz-2, and Protenix, predict assemblies containing proteins and small molecules^11–16^. More recent developments include OpenFold3, OpenDDE, ESMFold2, Protenix-v1 and Protenix-v2^17–21^. However, predicting MG ternary complexes remains challenging because both the protein–protein interface and the MG binding pose must be determined. These complexes can involve either two structured protein domains (Group 1) or a short sequence motif bound to a structured domain (Group 2)^3^. In addition, proteins brought together by an MG may lack the coevolutionary signals that help identify interfaces in naturally occurring complexes. In our previous benchmark, cofolding methods performed less accurately on MG complexes without closely related structures in their training data, with particularly poor performance on domain-domain interfaces^22^. These limitations motivate the development of methods specifically designed for MG ternary complex structure prediction.

Docking provides an alternative approach by assembling complexes from the structures of their components. Existing docking methods for PROTAC ternary complexes typically combine protein–protein docking, conformational sampling, and structural refinement, using the two protein-binding moieties and connecting linker to guide complex assembly^23–25^. Applying these strategies to MGs is challenging because their compact scaffolds do not provide the same two binding moieties and linker geometry. DeepTernary introduces the TernaryDB dataset and uses an SE(3)-equivariant network for ternary complex docking, with limited reported success^9^. Diffusion models have also been applied to several docking tasks, including protein–ligand docking with DiffDock^26^ and DiffBindFR^27^, protein–protein docking with DiffDock-PP^28^, and protein–peptide docking with DiffPepDock^29^. However, diffusion-based methods have not yet been specifically developed for MG ternary complex docking, which requires joint prediction of the relative protein arrangement and the MG binding pose.

Here we present DiffTernary, a diffusion model that jointly predicts protein arrangement and MG binding pose from component structures. To learn ternary interactions beyond the limited set of experimentally resolved MG complexes, we introduce Ternary-PDB, a collection of 76,702 ternary complexes with small-molecule or peptide bridges, assembled using structural filters and a release cutoff aligned with AF3. During training, one protein remains fixed while the partner protein and MG are jointly perturbed, allowing the model to learn their placement through denoising. We first evaluate the Tensor Product model trained on this dataset against AF3, OpenFold3, ESMFold2, Protenix-v1, Protenix-v2 and DeepTernary on the MG ternary complex benchmark MGBench^22^. With a single prediction, DiffTernary achieves a PPI prediction success rate comparable to AF3. DiffTernary ranks second with a best-of-25 PPI prediction success rate of 78.4%, behind ESMFold2 (90.9%) and ahead of AF3 (72.7%). We then train a separate Tensor Product model on TernaryDB for comparison with dedicated docking methods. Confidence-based selection among 40 samples yields acceptable PPI predictions for 41.5% of complexes, compared with 30.9% for DeepTernary. Finally, an E2Former variant^30^ increases training and sampling throughput 6.90-fold and 4.37-fold under the evaluated configurations. Because DiffTernary uses equivariant message passing without requiring multiple sequence alignments or Pairformer blocks, it provides a dedicated docking tool for future MG virtual screening and design.

## Results and Discussion

### Ternary-PDB establishes a structural data foundation for ternary interactions

To support training of DiffTernary, a dedicated ternary complex structure prediction model, we curated Ternary-PDB from experimentally resolved complexes with small-molecule or peptide bridges, thereby expanding the structural repertoire available for ternary systems (Fig. 1a). We constructed Ternary-PDB from Protein Data Bank (PDB) structures^31^ released on or before the AF3 training cutoff of 30 September 2021, allowing comparison with AF3 on subsequently released complexes. We included X-ray and cryo-EM structures at resolutions of 9 Å or better, following AF3’s limits on the number of chains per assembly and its structure preprocessing procedure^12^. We defined a ternary complex as two protein chains connected by a bridging molecule, with at least three heavy atoms of the molecule within 5 Å of each protein. We excluded bridging molecules that were covalently attached to either protein, contained metals, or were not resolved in the experimental density. In addition to small molecules, we included peptides shorter than 16 resolved residues that bridged two larger protein chains, allowing the dataset to cover both linear peptides and macrocycles. Each complex provides data on two proteins contacting a bridging molecule, the ternary geometry learned by DiffTernary. Lipid and cofactor bridges were therefore included alongside small molecules relevant to therapeutic MGs. Starting from 89,935 candidate PDB entries, 85,774 retained two protein chains after preprocessing, and 18,929 contained at least one qualifying bridging molecule. For comparison, TernaryDB was assembled from X-ray structures at 3.5 Å or better with R-free values no higher than 0.26 and did not apply AF3’s release cutoff^9^. Our collection of 76,702 ternary complexes represents a 3.4-fold increase in the number of complexes and a 4.6-fold increase in the number of PDB entries compared with TernaryDB, which contains 22,303 complexes^9^ (Fig. S1a).

**Figure 1.**
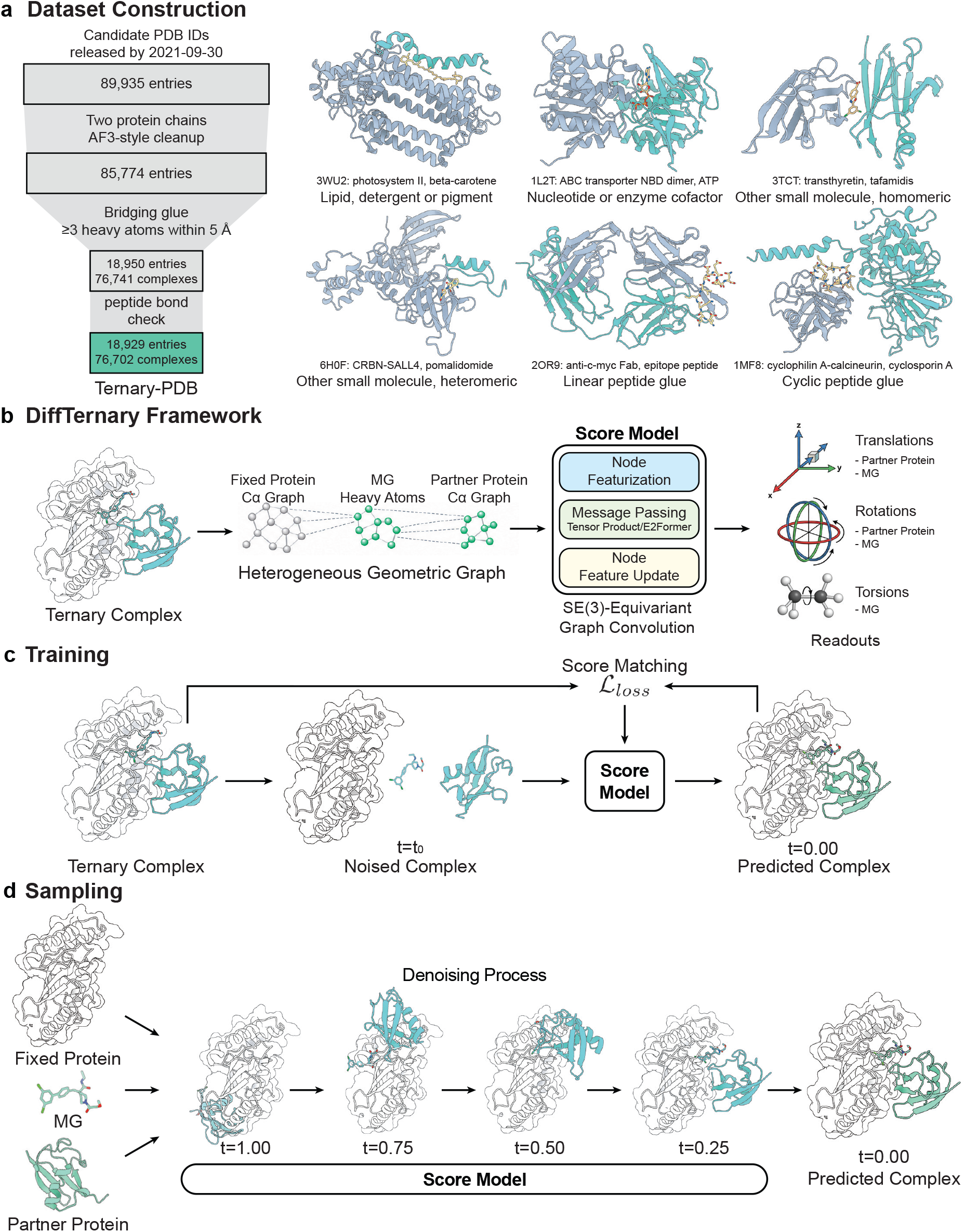
Overview of DiffTernary and Ternary-PDB. **a**, Dataset construction and examples of six bridge classes, with proteins in blue-gray and green and bridges as sticks. **b**, Graph of protein C*α* nodes and MG heavy atoms, with Tensor Product or E2Former message passing and translation, rotation and torsion score outputs. **c**, Score matching after joint perturbation of the movable protein and MG. **d**, Joint denoising from random component placements. In **b**–**d**, the fixed protein is white and the movable protein and MG are green.

Lipids, detergents and pigments were the most common bridges (39,114 complexes), followed by nucleotides and enzyme cofactors (15,029 complexes) (Fig. S1c). The dataset contains 153,404 protein-chain occurrences across its complexes. Source-organism annotations are available for 153,084 occurrences spanning 1,579 organisms. Photosynthetic species are prominent sources of pigment- and lipid-bridged complexes, alongside commonly studied model organisms. The 12 most represented organisms contribute approximately half of the annotated chain occurrences (Fig. S1d). Small molecules bridge proteins with different sequences in 4,198 complexes, the category most relevant to therapeutic MGs. Peptides bridge another 3,374 complexes, including 430 with cyclic peptides, which are absent from TernaryDB. Bridges with 20 to 55 heavy atoms account for 69.9% of complexes. Bridges with more than 100 heavy atoms occur in 1.2% of complexes and are mainly large lipids and pigments. Peptide bridges are generally larger, with a median of 76 heavy atoms (Fig. S1e).

We clustered the three components of each complex to control redundancy between training and validation. Protein chains were grouped by MMseqs2^32^ at 40% sequence identity, peptides shorter than 10 residues by identical sequence, and small-molecule bridges by chemical component identifier. Combining these component clusters yielded 23,827 complex clusters, twelve times the 1,982 clusters of TernaryDB. Of these, 13,412 contain a single complex (Fig. S1b). This expanded collection therefore includes additional combinations of protein sequence clusters and bridging molecules. MGBench^22^ contains 88 complexes with 68 MGs. Their protein sequences lie within the range represented in Ternary-PDB (Fig. S1g), and their molecular chemistries are similarly covered (Fig. S1f).

### Overview of DiffTernary

DiffTernary predicts an MG ternary complex from two protein conformations and the chemical structure of the MG. Both proteins are treated as rigid bodies, and the MG conformation changes through rotations around its rotatable bonds. One protein remains fixed while the partner protein and MG are docked jointly. Training includes both choices of fixed protein, and either protein can be fixed during inference. The heterogeneous graph contains one node per protein residue at its C_*α*_ atom and one node per MG heavy atom. The score network updates these features through equivariant message passing, implemented with Tensor Product convolutions or E2Former^30^. Both implementations predict translation and rotation scores for the partner protein and MG, together with MG torsion scores (Fig. 1b).

To model the coupled assembly of MG ternary complexes, we keep one protein fixed and jointly perturb the partner protein and MG during training. The network learns to reconstruct their relative placement by encoding interactions among all three components. The partner protein and MG are displaced from their native positions through rigid-body translations and rotations, while the MG is also perturbed along its rotatable bonds. Both movable components are perturbed at the same diffusion time, and the network learns the scores of these perturbations through denoising score matching (Fig. 1c). During inference, the partner protein and MG are initialized at random positions and orientations around the fixed protein, with MG torsion angles sampled uniformly and applied to an RDKit-generated conformer^33^. The network iteratively updates their positions and orientations and the MG torsion angles to generate a ternary structure (Fig. 1d). Cross-component edges are rebuilt at each denoising step using a distance cutoff that decreases over the diffusion process. The network therefore considers long-range interactions early in sampling and increasingly local interactions near the binding interface at later steps. A separate confidence model ranks candidates in the TernaryDB sampling experiments.

### DiffTernary achieves competitive PPI prediction against cofolding models

We evaluated the DiffTernary Tensor Product model trained on Ternary-PDB using MGBench, which comprises 88 experimentally determined MG ternary complexes released after the AF3 training cutoff of 30 September 2021^22^. Sequence comparisons with PDB structures released before that date divide the benchmark into 63 complexes with high sequence homology and 25 with low sequence homology. Comparisons included five cofolding methods, AF3, OpenFold3, ESMFold2, Protenix-v1 and Protenix-v2^12,17,19–21^, and the released DeepTernary model trained on TernaryDB^9^. The five cofolding models evaluated here used PDB training structures released on or before 30 September 2021. We applied the same cutoff when constructing Ternary-PDB, and all 88 MGBench complexes were released after this date. The docking methods use protein conformations and MG chemistry, whereas cofolding methods use protein sequences and MG chemistry. We evaluated PPI prediction accuracy using DockQ, which combines recovery of native protein contacts with deviations in the predicted protein arrangement^34^. DockQ > 0.23 corresponds to the CAPRI acceptable category^34,35^. A single blind prediction measures baseline docking accuracy, while oracle selection of the highest-DockQ pose among 25 samples assesses whether repeated sampling recovers an accurate structure and gives the accuracy attainable with ideal pose selection (Fig. 2a). With a single blind prediction per complex, DiffTernary achieved a PPI prediction success rate of 50.0%, compared with 51.1% for AF3. The best-of-25 PPI prediction success rates, defined by the highest-DockQ pose for each complex, reached 78.4% and 72.7%, respectively. ESMFold2 achieved the highest best-of-25 PPI prediction success rate at 90.9%. DiffTernary ranked second, ahead of Protenix-v2 (77.3%), OpenFold3 (59.1%) and Protenix-v1 (51.1%). DeepTernary produced acceptable PPI predictions for 38.6% of complexes with a single prediction and 43.2% with 25 samples, below DiffTernary in both settings despite potential training leakage from PDB entries shared between its training data and MGBench. On the 25 complexes with low sequence homology to PDB structures released before AF3’s cutoff^22^, DiffTernary produced acceptable PPI predictions for 40.0% of complexes with 25 samples, compared with 28.0% for AF3. The PPI prediction success rate increased with the sampling budget (Fig. S2).

**Figure 2.**
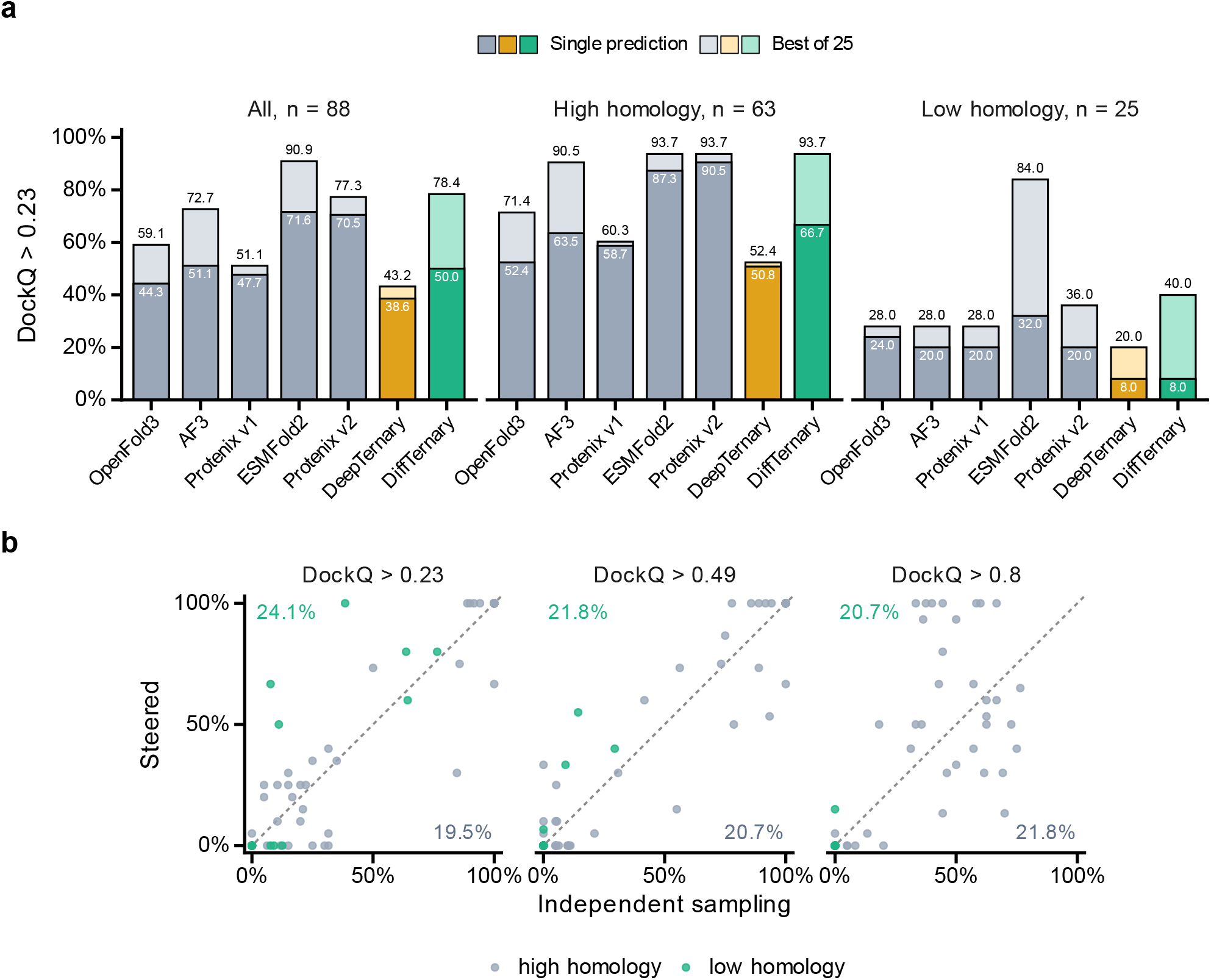
Prediction and steering on MGBench. **a**, Single-prediction and best-of-25 PPI prediction success rates above DockQ 0.23 on 88 complexes. Cofolding methods use five seeds with five samples each, and docking methods sample both choices of fixed protein. Scoring retains the highest DockQ across protein-chain copies. **b**, Each point represents one complex. The x and y coordinates are the percentages of 20 poses exceeding the indicated DockQ threshold under independent and Feynman–Kac sampling, respectively. Upper-left and lower-right labels give the percentages of all compared complexes with more and fewer successful poses under steering, respectively. Points on the diagonal have equal success counts under both methods.

We applied Feynman–Kac steering^36^ using a shape complementarity reward adapted from OpenDDE^18^. DockQ thresholds of 0.49 and 0.80 correspond to medium and high PPI prediction quality, respectively^34^. At the same budget of 20 poses per complex, steering increased the fraction of poses above DockQ 0.80 from 18.6% to 21.6% (Fig. 2b). On the low homology subset, the fractions above DockQ 0.23 and 0.49 increased from 11.6% to 17.5% and from 2.1% to 5.4%, respectively.

### DiffTernary outperforms docking methods on TernaryDB

For comparison in DeepTernary’s original benchmark setting, we trained a separate DiffTernary Tensor Product model on the TernaryDB MG training split. We evaluated both methods on the 94 MG test complexes^9^. We also included EquiDock^37^ and DiffDock-PP^28^ to assess the performance of protein–protein docking methods that do not take the MG as input. We first compared baseline docking accuracy using a single blind prediction from each method for each complex. EquiDock and DiffDock-PP achieved mean DockQ^34^ scores below 0.08 (Fig. 3a), with only a few predictions reaching the CAPRI acceptable threshold of DockQ > 0.23^35^ (Fig. 3b). DiffTernary achieved a mean DockQ of 0.306, compared with 0.228 for DeepTernary. DiffTernary also produced six times as many predictions with DockQ > 0.80 as DeepTernary, indicating better recovery of high-quality PPI predictions.

**Figure 3.**
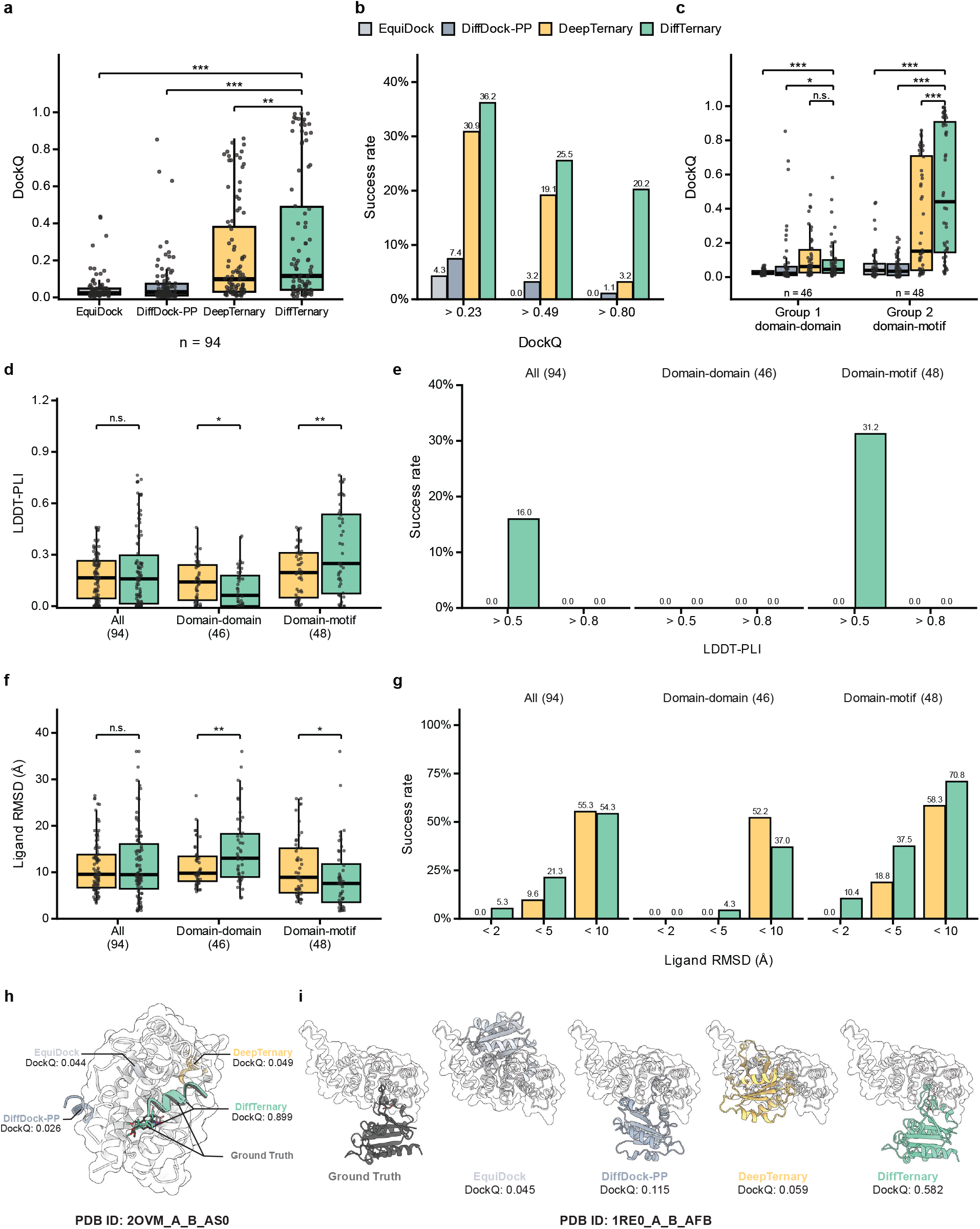
Single-prediction accuracy on TernaryDB. **a**, DockQ for four methods on 94 complexes. Boxes show the median and interquartile range, whiskers extend to 1.5 times that range, and points denote complexes. **b**, PPI prediction success rates at the indicated DockQ thresholds. **c**, DockQ for 46 domain-domain and 48 domain-motif complexes. **d, e**, MG LDDT-PLI and fractions above the indicated thresholds. **f, g**, Pocket-aligned MG RMSD and fractions below the indicated cutoffs. **h, i**, Predictions for 2OVM_A_B_AS0 and 1RE0_A_B_AFB, with the fixed protein in white and DockQ labels beside predictions. Two-sided paired Wilcoxon tests compare DiffTernary with the other methods. Holm correction applies within each comparison family in **a, c.** Symbols denote ^*^*P* < 0.05, ^**^*P* < 0.01 and ^***^*P* < 0.001. The abbreviation n.s. denotes a non-significant difference.

We next compared performance across the two types of MG ternary complex defined by Rui et al.^3^, Group 1, in which two structured protein domains interact, and Group 2, in which a short sequence motif interacts with a structured domain. Both DiffTernary and DeepTernary performed better on Group 2 than on Group 1 (Fig. 3c), consistent with the DeepTernary study^9^. For Group 1, few predictions achieved DockQ > 0.49, and none of the methods accurately recovered the MG binding pose. The improvement of DiffTernary over DeepTernary was mainly observed in Group 2, where DiffTernary achieved acceptable DockQ scores for more than half of the complexes and recovered substantially more structures with DockQ > 0.80. We also evaluated MG placement using the local distance difference test for protein–ligand interactions (LDDT-PLI) (Fig. 3d). DiffTernary was the only method to achieve LDDT-PLI scores above 0.5 (Fig. 3e). Pocket-aligned ligand root-mean-square deviation (RMSD) provided a complementary measure of MG placement (Fig. 3f). DiffTernary was also the only method to predict MG poses within 2 Å of the experimental structures (Fig. 3g). It also approximately doubled the fraction of MG poses with ligand RMSD below 5 Å compared with DeepTernary. These improvements were primarily observed in Group 2, where DiffTernary achieved better median ligand RMSD and LDDT-PLI scores.

Replacing each MG with a single carbon atom at its centroid reduced mean DockQ from 0.306 to 0.273 and the PPI prediction success rate from 36.2% to 29.8%, with both protein conformations unchanged (Fig. S3). The effect was strongest for Group 1 domain-domain complexes, where the PPI prediction success rate fell from 10.9% to 4.3%, supporting a contribution of MG chemical information to protein arrangement. We also selected representative examples for visualization. For 2OVM_A_B_AS0, a Group 2 complex, DiffTernary accurately reproduced the experimental protein arrangement, whereas the other three methods placed the partner protein incorrectly (Fig. 3h). For 1RE0_A_B_AFB, a Group 1 complex, DiffTernary was the only method to produce an acceptable PPI prediction (Fig. 3i). These results demonstrate that DiffTernary improves the prediction of MG-mediated PPIs compared with existing docking methods.

### Repeated sampling and confidence ranking improve docking accuracy

We then examined how repeated sampling and pose selection affect the docking accuracy of DiffTernary trained on TernaryDB. We next assessed whether repeated sampling could recover accurate poses missed by a single prediction. For each complex, we generated up to 40 samples and used the native structure to identify the most accurate pose, giving an oracle estimate of the accuracy attainable with ideal pose selection. On the 94 MG complexes, the PPI prediction success rate (DockQ > 0.23) increased from 24.6% with one sample to 52.1% with the best of 40 samples (Fig. 4a). By comparison, DeepTernary improved by less than two percentage points across 40 runs, whereas EquiDock produced the same prediction in every run. DiffDock-PP also benefited from repeated sampling, reaching 48.9% with 40 samples. DiffTernary showed similar improvements at the higher DockQ threshold of 0.80 (Fig. 4b). For MG placement, 16.0% of complexes had at least one predicted MG pose within 2 Å of the experimental structure among 40 samples, whereas DeepTernary did not achieve this accuracy for any complex (Fig. 4c).

**Figure 4.**
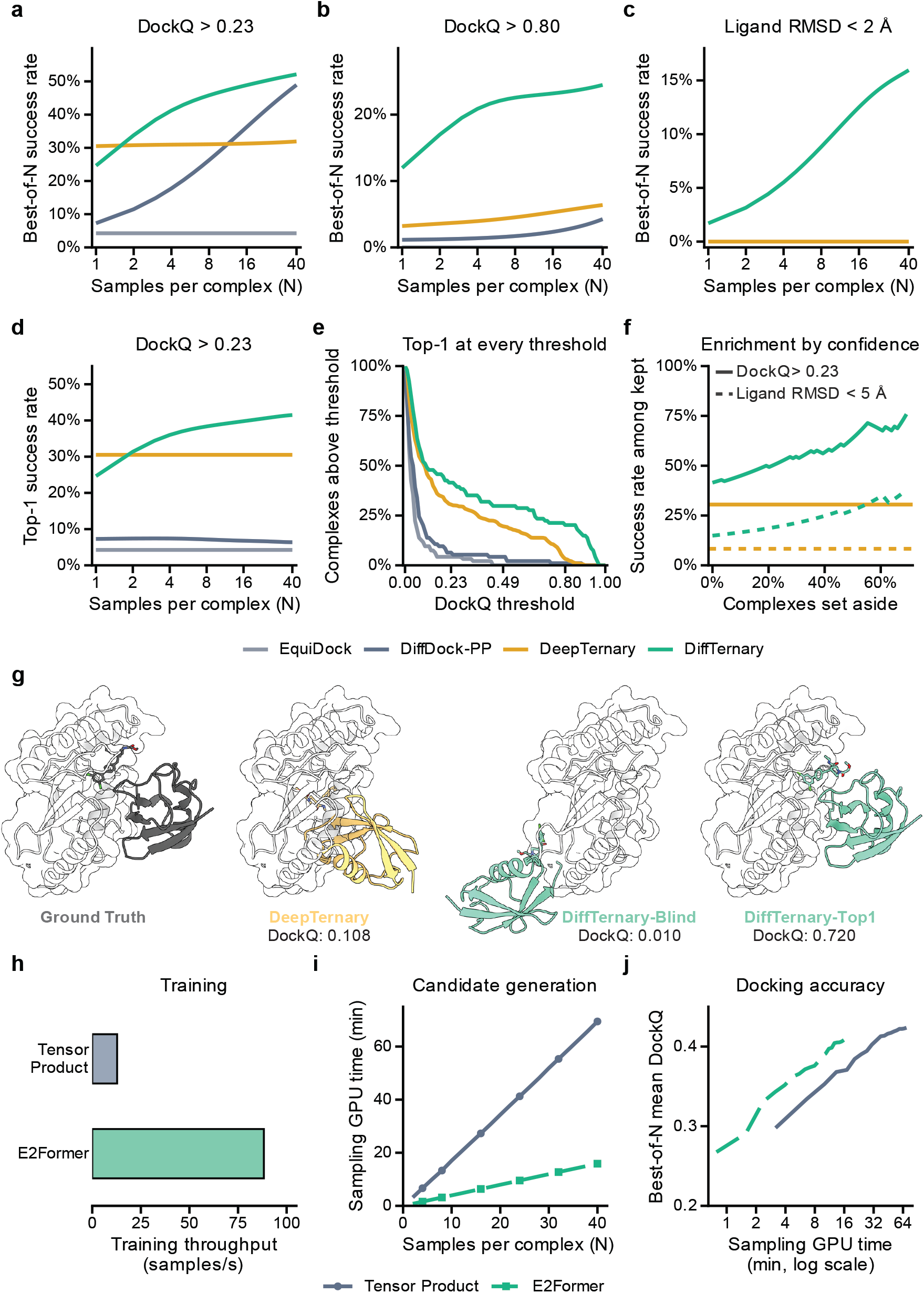
Sampling, ranking and efficiency on TernaryDB. **a, b**, Best-of-*N* PPI prediction success rates at DockQ thresholds of 0.23 and 0.80. **c**, Best-of-*N* MG recovery within 2 Å. **d, e**, Returned-pose PPI prediction success rates against *N* and at *N* = 40. DiffTernary and DiffDock-PP use confidence selection. Values in **a**–**d** average over subsets of 40 samples per complex. **f**, PPI and MG prediction success rates after confidence filtering. Horizontal lines give expected PPI and MG prediction success rates for a single DeepTernary prediction. **g**, Native and predicted 7M2K_A_B_GZM structures, with the fixed protein in white. **h**, Training throughput on eight H800 GPUs using FP32 and per-GPU batch sizes of 1 for Tensor Product and 16 for E2Former. **i**, Sampling time against *N*. **j**, Mean oracle DockQ against sampling time on a logarithmic axis.

Oracle selection requires the native structure. We trained a separate confidence model to rank the generated poses for selection at inference. The confidence model was trained on DiffTernary predictions to rank their protein–protein interfaces according to interface RMSD (eq. (8)). With 40 samples per complex, selecting the highest-scoring pose increased the PPI prediction success rate to 41.5% (Fig. 4d), whereas the ranking model of DiffDock-PP showed little improvement with additional samples. DiffTernary also outperformed DeepTernary across DockQ thresholds (Fig. 4e).

We also tested whether confidence scores identify reliable predictions across different complexes, an evaluation termed selective prediction^26^. After excluding the approximately 70% of predictions with the lowest confidence scores, the PPI prediction success rate among the remaining complexes increased from 41.5% to 78.6%. The fraction of MG poses with ligand RMSD below 5 Å also increased from 14.9% to 39.3%. The corresponding rates for a DeepTernary prediction taken at random from its 40 runs were 30.9% and 8.3%, respectively (Fig. 4f). The complex 7M2K illustrates the benefit of sampling and confidence ranking (Fig. 4g). A single DiffTernary sample placed the partner protein incorrectly (DockQ 0.010), whereas DeepTernary achieved a DockQ of 0.108. By sampling 40 poses and selecting the highest-scoring one, DiffTernary produced a substantially more accurate PPI prediction, at DockQ 0.720.

### E2Former accelerates DiffTernary training and sampling

Finally, we investigated whether E2Former could accelerate training and repeated sampling of DiffTernary trained on TernaryDB. We implemented an E2Former variant that computes geometric tensor products at nodes and reuses them across edges during message passing^30^. The variant retains the graph construction, diffusion variables and training objectives of the Tensor Product model. On eight H800 GPUs with FP32 precision, measured training throughput was 12.79 samples per second for Tensor Product and 88.29 for E2Former, a 6.90-fold increase. The per-GPU batch sizes were 1 and 16, respectively (Fig. 4h, Table S1). We compared candidate generation in a separate timed sampling experiment on the same 94 TernaryDB test complexes.

Each model generated 20 candidates under each choice of fixed protein, giving 40 candidates per complex. On one A800 GPU, Tensor Product required 4,167.14 s and E2Former required 952.71 s, corresponding to a 4.37-fold increase in candidate generation throughput (Fig. 4i). Tensor Product used FP32 precision and batch size 1, while E2Former used BF16 and batch size 16. In this experiment, the blind predictions achieved mean DockQ values of 0.2577 for Tensor Product and 0.2358 for E2Former. Selecting the best of 40 candidates increased these values to 0.4237 and 0.4082, with acceptable PPI predictions obtained for 56.4% and 50.0% of complexes, respectively. Mean oracle DockQ first exceeded 0.40 after 12.7 min with E2Former and 30.7 min with Tensor Product (Fig. 4j). The time required to accumulate these candidate sets and their resulting PPI prediction accuracy are summarized in Table S1. E2Former increased sampling throughput while retaining a similar mean oracle DockQ in this comparison.

## Discussion

DiffTernary and Ternary-PDB establish a modeling framework and a structural data foundation for MG ternary complex prediction. Learning from a broad collection of experimentally resolved ternary assemblies allows the model to draw on interactions beyond the relatively small set of known MG complexes. Joint denoising couples the placement of the MG and partner protein around a fixed protein, allowing their relative positions to be updated through message passing among all three components. On MGBench, DiffTernary achieved blind PPI prediction accuracy comparable to AF3 and higher oracle accuracy than AF3, OpenFold3 and both Protenix versions. A separate model trained on TernaryDB outperformed DeepTernary on its MGD test set^9^. These results support the use of component structures and learned ternary interactions for MG docking. The combination of a dedicated docking model and an expanded structural dataset provides a basis for studying how small molecules bring protein partners into compatible binding arrangements.

Repeated sampling makes exploration of alternative ternary arrangements a central part of prediction. Building on diffusion-based docking^26,38^, DiffTernary generates candidate structures that differ in protein placement and MG binding pose. The increase in PPI prediction success under oracle selection shows that additional samples recover accurate structures across more complexes. Confidence ranking selects among completed predictions, while Feynman–Kac steering uses geometric information during generation^36^. The shape complementarity term adapted from OpenDDE connects steering to the fit between interacting molecular surfaces^18^. E2Former addresses the computational cost of this exploration through more efficient equivariant message passing^30^. Its increased throughput, accompanied by similar mean oracle DockQ in the timed comparison, makes larger candidate sets accessible within a given computational budget. Together, these findings connect prediction accuracy with the practical ability to explore and assess alternative binding arrangements.

The biological relevance of MG recognition makes structural prediction valuable for both mechanistic studies and molecular design^1,3^. Predicted ternary arrangements can provide testable hypotheses about how an MG engages its protein partners and which contacts may contribute to complex formation. DiffTernary offers a dedicated computational approach to this task without requiring multiple sequence alignments or Pairformer computation during docking. Its sampling efficiency and the structural coverage of Ternary-PDB provide complementary foundations for evaluating candidate MG complexes and organizing structural evidence across systems. As experimental structures and measurements of MG activity accumulate, successive cycles of computational prediction and experimental evaluation could continuously refine the relationship between ternary geometry and biological function. This interaction between modeling and experiment offers a broader route toward systematic MG discovery and design.

## Methods

### Ternary-PDB data collection and filtering

Ternary-PDB was assembled from the Protein Data Bank^31^ using the structure eligibility and preprocessing rules of AF3^12^ (Fig. 1a). Candidate entries satisfied the three target filters in AF3 Supplementary Information (SI) 2.5.4^12^. They were released on or before 30 September 2021, had resolution 9 Å or better, and contained at most 300 polymer chains. Each entry also had to contain at least one protein entity. Two classes of entry were retrieved. Entries containing a non-polymer entity required at least two polymer chains in the assembly. Entries without non-polymer entities required at least three polymer chains, allowing a peptide to act as the bridge. Only the first biological assembly was used, following the assembly expansion used for AF3 training (SI 2.1). The search returned 89,935 entries.

Assemblies were processed using the AF3 parsing rules (SI 2.1). Hydrogens and waters were removed, alternative locations were resolved by highest occupancy, and selenomethionine was converted to methionine. Modified protein residues were mapped to their parent amino acids and retained only the parent’s atoms, using the SCOP mapping adopted for AF3 clustering (SI 2.5.3). The filters in SI 2.5.4 were then applied. Polymer chains composed entirely of unknown residues were removed, as were protein chains with fewer than four resolved residues or with sequence-consecutive C_*α*_ atoms separated by more than 10 Å. Units were removed when more than 30% of their atoms lay within 1.7 Å of another unit. Mutual clashes were resolved in favor of the chain with the smaller clashing fraction. The crystallization aids in SI Table 9 were removed from crystallographic structures. Ions, lipids, glycans and cofactors were retained. At least two protein chains survived in 85,774 entries.

A ternary complex comprised two protein chains and one bridging molecule with at least three heavy atoms within 5 Å of each protein. Bridges annotated by wwPDB as covalently bonded or disulfide-linked to a protein were excluded to retain molecules free to move. Metal-containing bridges were excluded because they could not be processed by our featurization pipeline. Bridges with at least half their heavy atoms at zero occupancy were excluded because they were insufficiently resolved in the density. Crystallization aids, excluded ligands, glycans and ions listed in AF3 SI Tables 9–12 were excluded as bridges. Because the bridge defines the ternary complex, this selection applies AF3’s exclusions to the molecule around which it is assembled. A peptide bridge was a chain of fewer than 16 resolved residues contacting two protein partners of at least 16 residues each. It had to be gapless and contain no unknown residues. Modified residues and terminal caps were retained as deposited, consistent with treating the peptide bridge as a molecule. When one bridging molecule contacted the same protein pair at multiple sites, the site with the most contacts was retained. Symmetry copies within each assembly were counted once. These steps yielded 76,741 complexes from 18,950 entries.

Peptide bridges were required to form a connected molecule with consecutive residues joined by C–N bonds. Every such bond had to be between 1.1 and 2.2 Å, with at most one bond per residue pair and no interatomic distance below 1.0 Å. These checks excluded 39 complexes. The resulting dataset, Ternary-PDB, contains 76,702 complexes from 18,929 entries, of which 3,374 are bridged by a peptide and 430 of those by a cyclic peptide.

### Ternary-PDB clustering and dataset splitting

Chains were clustered following AF3 SI 2.5.3. Protein chains were grouped with MMseqs2^32^ at 40% sequence identity, peptides shorter than 10 residues by identical sequence, and small-molecule bridges by chemical component identifier. Each chain was represented by its full entity sequence, with modified residues mapped to their parents. This produced 13,216 chain clusters. AF3 defines an interface cluster by the clusters of its two constituent chains. We extended this definition to a ternary complex by combining the unordered pair of protein clusters with the bridge cluster. The 76,702 complexes formed 23,827 complex clusters, of which 13,412 contained a single complex.

Complexes sharing a complex cluster or a PDB entry were grouped together. We excluded 172 complexes whose protein chain-length product exceeded 1.5 million, as this product determines the number of possible edges between the proteins. The remaining groups were shuffled with random seed 0, and 150 groups containing a single complex each were selected for validation. This gave 76,380 training complexes and 150 validation complexes. The two splits share no complex clusters or PDB entries, although individual protein sequence clusters can occur in both. MGBench served as the external test set.

### TernaryDB and the MG split

The DeepTernary comparison used TernaryDB, the ternary complex collection released with that method^9^. Each entry is a crystallographically resolved complex reduced to the two proteins forming the induced interface and the bridging molecule. Entries are identified by the PDB code, two author chain identifiers and the MG chemical component identifier. In the released MG split, the 94 test complexes belong to 44 sequence clusters containing 276 complexes in total. The other 182 cluster members are withheld from training, and 137 form the validation set used for checkpoint selection. The 22,046 training complexes belong to 1,982 clusters, which define the training sampler weights. Of these, 882 could not be parsed or used owing to problems in the deposited data. Each of the remaining 21,164 complexes was loaded twice with the two bound protein conformations and MG, once under each choice of fixed protein, giving 42,328 training samples per epoch.

### Pose representation and diffusion

Each protein is represented by one node per residue at its C_*α*_ atom and the MG by one node per heavy atom. With protein *P*_1_ fixed, a structure is specified by 12 + *b* pose parameters. These comprise translations and rotations of the partner protein *P*_2_ and MG *L*, and one dihedral angle for each of the *b* rotatable MG bonds. Bond lengths, bond angles and ring geometry are fixed to an RDKit-generated conformer^33^. The map *A* converts pose parameters into coordinates by translating each movable body, rotating it about its centroid and adjusting each MG dihedral through rotation of the atoms on one side of its bond. The MG is then realigned to its rigidly transformed reference, preserving its centroid and making the choice of bond side immaterial. On the structures associated with the given conformer, this map is invertible and *A*^−1^ recovers the pose parameters. We use *R*(*u*) for a rotation of angle ∥*u* ∥ about axis *u*. Fixing the bound conformation of each protein while predicting their relative placement is the rigid-body approximation used in most protein docking^39^. DiffTernary learns *p*(*P*_2_, *L*| *P*_1_) by denoising diffusion over these parameters, preserving the component geometry throughout the sampled rearrangements.

The pose space comprises five channels, two translations in R^3^, two rotations in SO(3) and *b* torsion angles on the circle.

The noise scales follow a geometric schedule,

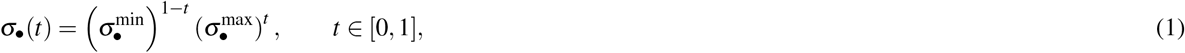

with *σ*_tr_ ∈ [0.1, 30] Å, *σ*_rot_ ∈ [0.03, 1.55] and *σ*_tor_ ∈ [0.0314, 3.14]. Translations are perturbed by isotropic Gaussians, rotations by the isotropic Gaussian distribution on SO(3), whose rotation angle *ω* has density

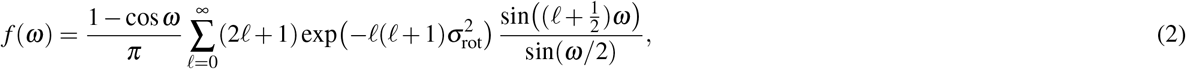

and torsions by wrapped normal distributions on the torus^40–42^. The series in eq. (2) and the corresponding scores are tabulated once and interpolated during training^26,28^. A single diffusion time *t U* (0, 1) is sampled per complex and shared by all five channels, while the perturbations themselves are sampled independently within each channel. The network is trained by denoising score matching, with each channel normalized by its typical score magnitude at the sampled noise level to balance contributions across *t*. The five terms are weighted 0.167, 0.167 and 0.166 for the translation, rotation and torsion of the MG, and 0.25 each for the translation and rotation of the partner protein.

### Score model

The score network uses a heterogeneous graph of residues from both proteins and MG atoms, with SE(3)-equivariant tensor-product convolutions^26,28^ implemented in e3nn^43^. Each MG atom has sixteen categorical features describing its element, chirality, degree, formal charge, implicit valence, attached hydrogens, radical electrons, hybridization, aromaticity, ring count and membership of rings of sizes three through eight. Protein residues carry an amino-acid type from a vocabulary of thirty-eight entries, comprising the twenty standard residues, seventeen modified residues and a catch-all category, together with a 1280-dimensional ESM2-650M embedding^44^. Covalent MG edges encode single, double, triple and aromatic bond orders as one-hot features. Separate embedding tables map each categorical feature to the scalar node and edge inputs used in eqs. (3) and (4). MG bond orders are assigned from wwPDB chemical component definitions^31^, and stereochemistry is assigned from the deposited coordinates. Only single bonds outside rings are treated as rotatable. Protein edges connect residues within 30 Å, with at most 24 neighbors per residue. MG edges combine covalent bonds with a 5 Å radius graph.

The edges between the three components are rebuilt at every denoising step with a cutoff that follows the current noise level, 3*σ*_tr_(*t*) + 20 Å between MG and protein and 3*σ*_tr_(*t*) + 40 Å between the two proteins. The larger early cutoffs support interactions between separated components, and the neighborhood contracts as denoising proceeds. Edges form four groups connecting MG atoms, residues within each protein, MG atoms to protein residues, and residues across the two proteins. Each convolution layer has a separate set of tensor-product weights for each group. Node features are first refined within each component separately by one embedding layer, then processed jointly by four interaction layers.

A node *i* carries a position *X*_*i*_ and a feature *h*_*i*_, where *h*_*i*_ is a direct sum of irreducible representations of SO(3) with 32 scalar channels and 6 vector channels. The second layer adds the parity-even vector block, and the third adds the parity-odd scalar block. Parity describes how a feature transforms under spatial inversion, *X* ↦−*X* . Even features remain unchanged, while odd features change sign. Write *r*_*i j*_ = *X*_*j*_ − *X*_*i*_ for the vector along an edge, *Y* for the real spherical harmonics up to ℓ = 2, *µ* for a Gaussian radial expansion, *η*(*t*) for the sinusoidal embedding of the diffusion time, *N*_*i*_ for the neighbors of *i* and *k*(*i j*) ∈ {1,…, 4} for the group an edge belongs to. One layer of the network is

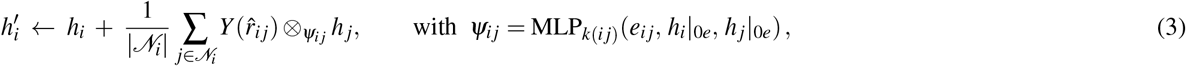

where ⊗ _*ψ*_ is a fully connected tensor product whose weights are produced for each edge by the network MLP_*k*_ of that group, *h* | _0*e*_ denotes the scalar channels of a feature and the residual term is zero-padded to the width of the layer output. Empty neighborhoods contribute zero messages. The edge feature itself is fixed over the layer stack,

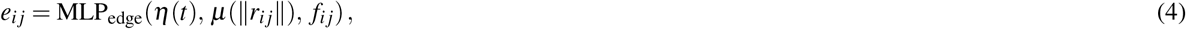

where *f*_*i j*_ contains bond features for covalent MG edges and is empty for other edges. Separate MLP_edge_ networks encode MG edges, protein edges and edges between components. The tensor-product weights *ψ*_*i j*_ are specific to the four relation groups. The tensor product couples node representations with the spherical harmonics of the edge direction, which makes each layer equivariant to rotations of the whole complex. The four groups are the MG to itself, each protein to itself, the MG to either protein and the two proteins to each other.

Two output heads predict the five score components from the final interaction features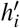. For a movable body *B*, either the partner protein *P*_2_ or the MG *L*, an additional set of edges runs from every node of *B* to the centroid 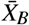 of that body, and one tensor-product convolution over them gives two parity-odd and two parity-even vectors,

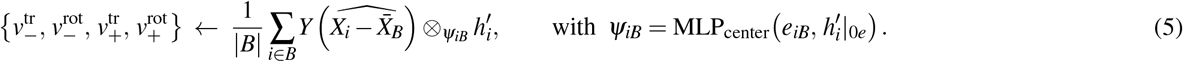

The edge features *e*_*iB*_ are constructed as in eq. (4). The two translation vectors and the two rotation vectors are summed separately. A small network then maps each vector magnitude and the diffusion time to an output magnitude while preserving its direction,

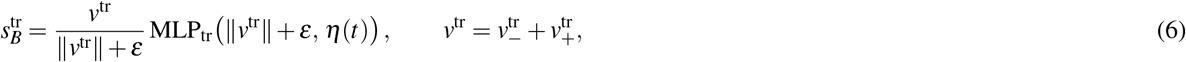

where *ε* = 10^−6^, and likewise 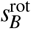 from *v*^rot^. The partner protein and the MG share the weights of both, since they undergo the ^42^ same rigid-body diffusion. The torsion head follows the pseudotorque construction of Jing et al. . For a rotatable bond *g* of the MG, with center *X*_*g*_ and axis *r*_*g*_, the MG atoms within 5 Å of *X*_*g*_ send messages whose spherical harmonics are coupled to the ℓ = 2 harmonic of the axis,

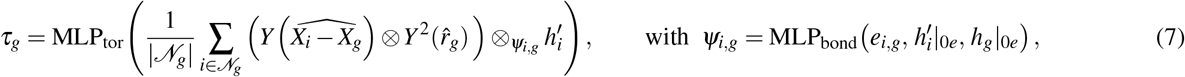

where *e*_*i,g*_ is constructed as in eq. (4), *h*_*g*_ is the sum of the features of the two bonded atoms, and MLP_tor_ reads the parity-odd and parity-even scalars of the message through two layers with no bias to produce the torsion score *τ*_*g*_. The five score outputs 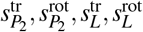 and *s*^tor^ = (*τ*_1_, …, *τ*_*b*_) are finally rescaled to match the magnitude of the true score at that noise level, by 1/*σ*_tr_ for the translations and by the tabulated score normalization factors *ν*_rot_(*σ*_rot_) and *ν*_tor_(*σ*_tor_) of the IGSO(3) and wrapped normal kernels for the rotations and the torsions.

### Confidence model

Samples generated for one complex are ranked by a separate confidence network, which reuses the score architecture up to the final interaction layer, pools the MG and partner features and emits one scalar per pose. Training optimizes the ordering of poses generated for the same complex. Writing *ϕ*_*i*_ for the score of pose *i* and *ρ*_*i*_ for the RMSD of the protein–protein interface it forms, the loss runs over the ordered pairs of one complex whose interfaces differ by more than a margin *m*,

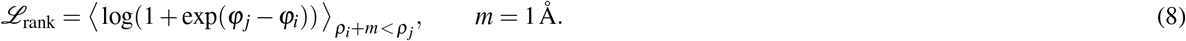

The loss encourages the pose with the lower interface RMSD to receive the higher score. Pairs within the margin are omitted. Batches are assembled by complex, so that the poses compared always belong to the same structure.

### Training and inference

Settings of the Tensor Product score and confidence networks are given in Table S2. Training and sampling are described in Algorithms 1 and 2. In Algorithm 1, each residual is divided by the typical score magnitude at its noise level. This is *ν*_tr_ = 1/*σ*_tr_ for translation and the tabulated score normalization factors *ν*_rot_(*σ*_rot_) and *ν*_tor_(*σ*_tor_) for rotation and torsion. The brackets ⟨·⟩ average over the components of each term. The TernaryDB score model was trained on eight 80 GB H800 GPUs for 52 epochs over 49 h. Validation DockQ was evaluated every five epochs on 137 complexes. The reported epoch-42 weights are close to the validation peak at epoch 40 (Fig. S4). From epoch 31 onward, the validation score matching loss exceeds the zero-prediction baseline, while validation DockQ and pose errors remain stable or continue to improve. Checkpoint selection therefore used validation DockQ. The confidence model was trained on four of the same GPUs for 2.8 h.

#### Algorithm 1

DiffTernary training, one epoch

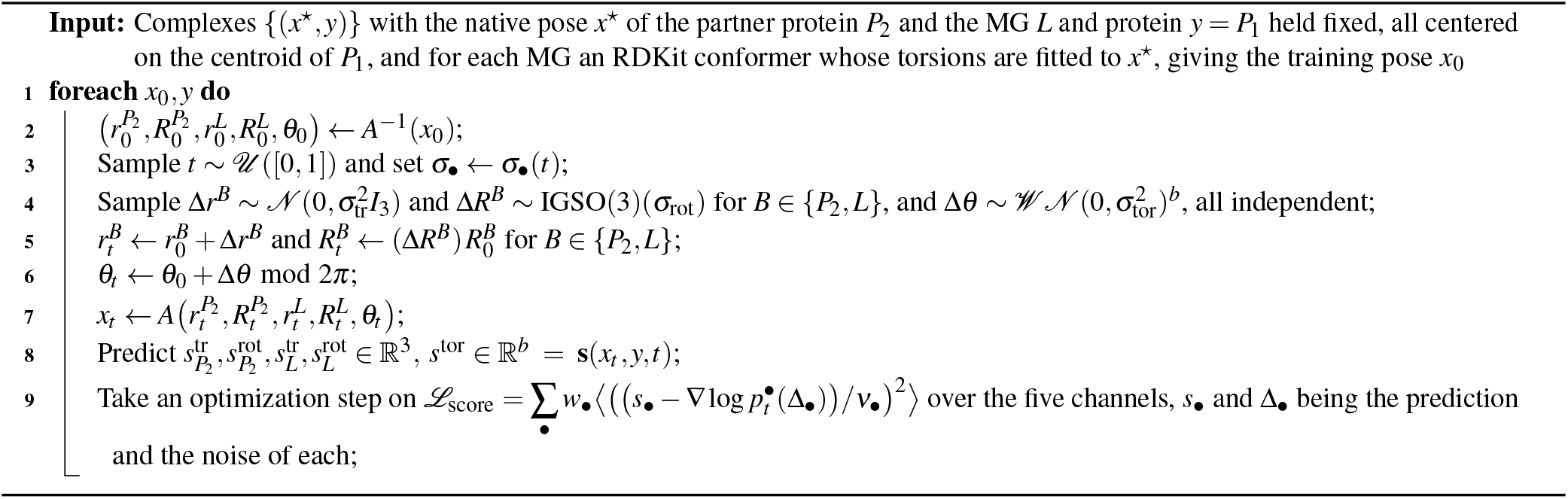

#### Algorithm 2

DiffTernary inference, one sample

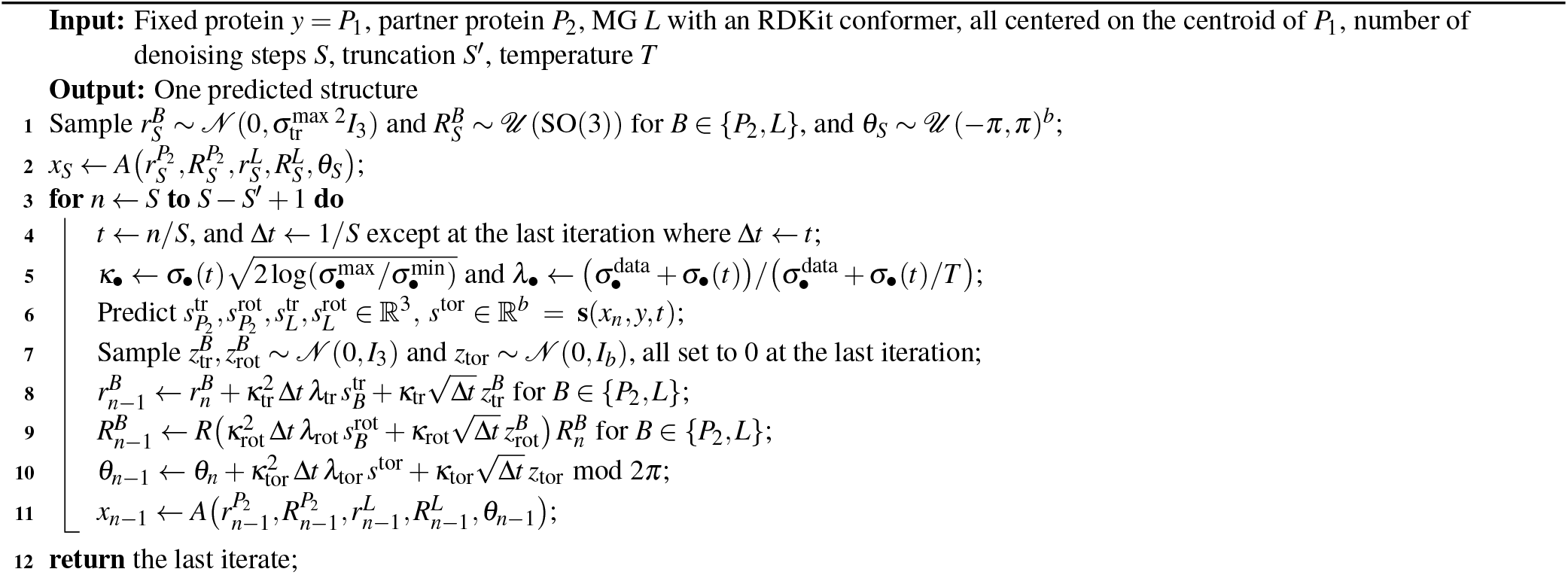

The Tensor Product score model for Ternary-PDB was trained on four 80 GB H800 GPUs, with checkpoint selection based on validation DockQ. Each epoch comprised 42,328 samples under cluster weighting, matching one pass over TernaryDB and providing a common sample-count scale for the two training runs. Validation DockQ was evaluated every five epochs on 150 complexes under both choices of fixed protein. The reported ep116 snapshot has epoch index 115. Loss measurements extend through epoch 117, and docking evaluations through epoch 115, where the highest recorded mean DockQ was 0.4999. Of the 299 valid predictions, 72.2% exceeded DockQ 0.23 (Fig. S5). After initialization, training and validation losses generally remained below the zero-prediction baseline, with isolated spikes and one non-finite validation value described in the figure caption.

The Tensor Product score models were trained with Adam at a learning rate of 10^-3^ and no weight decay. An exponential moving average of the weights (decay 0.999) was used for validation and saved checkpoints, and the learning rate followed a plateau scheduler based on validation loss. Training complexes were sampled with replacement under weights giving every sequence cluster the same total mass, so that large families of near-duplicate structures do not dominate the gradient. Checkpoints were selected on validation DockQ. At inference, each complex was denoised from a random placement of the partner protein and the MG using 20 time points and 19 updates. The final update advances to *t* = 0 without added noise. Samples were generated with the low-temperature scheme of Corso et al.^38^, in which the predicted score of each channel is multiplied by *λ* = (*σ*_data_ + *σ* )/(*σ*_data_ + *σ* /*T* ) before the update while the noise term is left alone, with *σ*_data_ the geometric mean of the noise range of that channel. For *T* > 1, *λ* > 1 increases the drift relative to the unchanged noise term. The resulting sharpening of the sampled distribution gives the low-temperature scheme its name. We used *T* = 4 and left the churn term of the scheme at zero, which puts *λ* between 1.04 and 3.44 over the noise range of the translations, between 1.10 and 2.93 for the rotations and between 1.07 and 3.14 for the torsions. Fig. S6 shows the sensitivity to the temperature and to the number of denoising steps.

### Steered sampling

Feynman–Kac steering favors trajectories with increasing reward^36^. For each complex and each choice of fixed protein, we ran two independent groups of five particles. Every two denoising steps, particles within each group were resampled using systematic resampling with weights *w*_*i*_ ∝ exp(Δ*r*_*i*_/*τ*), where Δ*r*_*i*_ is the change in reward since the previous resampling step.

The first resampling step used equal weights. At subsequent steps, *τ* was the standard deviation of the reward changes within the group, with a minimum value of 10^-9^. This procedure produced 20 final poses per complex across the two assignments. The unsteered control used the first ten sampling rounds from each assignment, with the same model weights and sampling temperature.

The reward measures shape complementarity in the rigid-body denoised estimate at each step, following the geometric construction of OpenDDE^18^. Let *A* contain the fixed protein and MG and *ℬ* contain the partner protein. Each protein residue is represented by its C_*α*_ atom and each MG heavy atom by one point. For a point *u* at *c*_*u*_, the local outward direction is estimated from the Gaussian density of points on the same side,

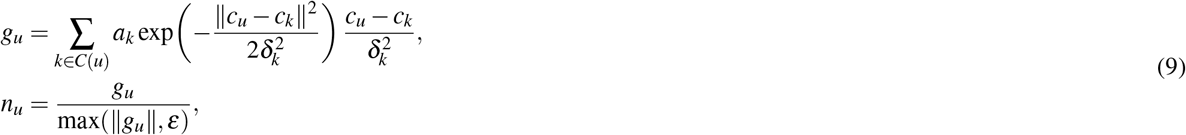

where *C*(*u*) is *A* or *ℬ* according to the side containing *u*, and *ε* = 10^−6^. Protein points use density width *δ*_*k*_ = 4 Å and weight *a*_*k*_ = 1, while MG atoms use *δ*_*k*_ = 2 Å and *a*_*k*_ = 0.125. For *u* ∈ *A* and *v* ∈ *ℬ*, define *d*_*uv*_ = ∥*c*_*v*_ − *c*_*u*_∥ and 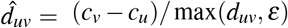. The four geometric factors are

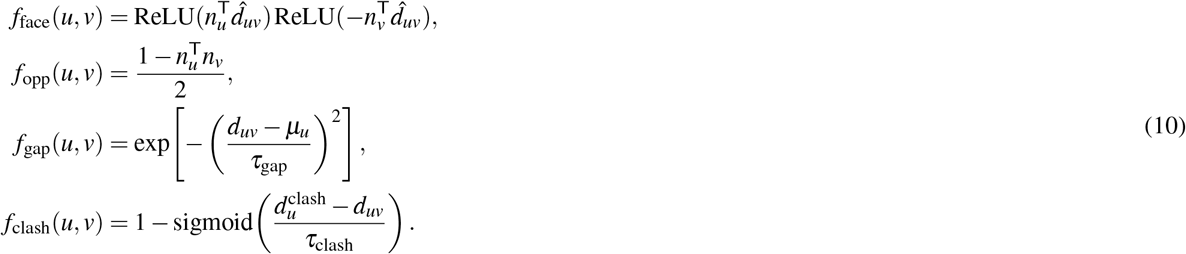

The facing term favors points whose outward directions face one another, and the opposing-normal term favors complementary surface orientations. The gap term favors an appropriate separation, and the clash term smoothly penalizes close contacts. Their product gives the pair score,

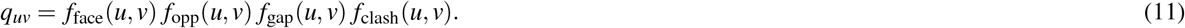

For pairs of protein points, *µ*_*u*_ = 6.5 Å and 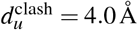. For an MG atom paired with a partner protein point, these values are 4.5 Å and 3.0 Å, respectively. Both pair types use *τ*_gap_ = 2.5 Å and *τ*_clash_ = 0.5 Å.

A pair is valid when *d*_*uv*_ ≤ 12 Å and both density-gradient magnitudes exceed 10^-3^. Writing *V*_*v*_ for the valid points in *A* paired with partner point *v*, pair scores are pooled using distance weights,

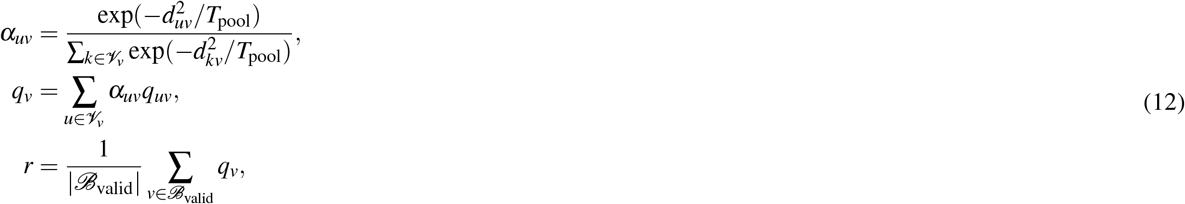

where *T*_pool_ = 25 Å^2^ and*ℬ* _valid_ contains partner points with at least one valid pair. The reward is zero when this set is empty. The geometric factors and their weighted averages lie in [0, 1], and the reward has no learned parameters. We use *r* from eq. (12) to compute the reward increments for resampling. To evaluate steering, we calculate the fraction of poses above each DockQ threshold for each complex and average these fractions across complexes, estimating the success probability of a single pose from steered sampling.

### Evaluation

Predictions were evaluated against experimental structures using the three metrics employed in the cofolding benchmark of MG ternary complexes^22^. DockQ assesses PPI prediction accuracy^34^, LDDT-PLI measures preservation of contacts between the MG and its binding pocket, and ligand RMSD measures the MG pose after binding-site superposition^45^. The first of the three is

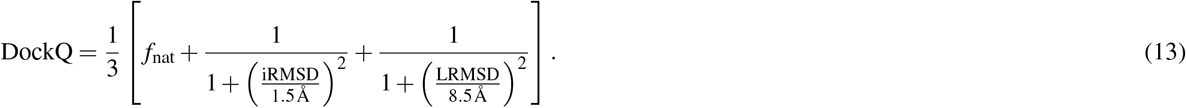

Here, *f*_nat_ is the fraction of native contacts recovered, iRMSD is the backbone RMSD after interface superposition, and LRMSD is the backbone RMSD of the shorter protein after superposing the longer protein. The second is

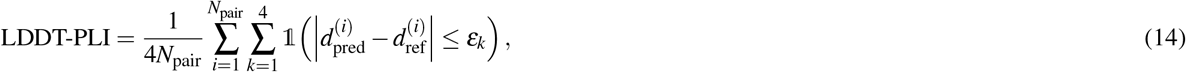

where the sum runs over the *N*_pair_ pairs consisting of one MG atom and one protein atom within 6 Å of each other in the native structure, *d*^(*i*)^ is the separation within pair *i* measured in the predicted and in the native structure, 1 is the indicator function and the tolerances *ε*_*k*_ are 0.5, 1.0, 2.0 and 4.0 Å. The third is

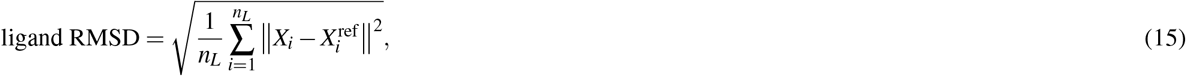

with *n*_*L*_ the number of heavy atoms of the MG and *X*_*i*_ and 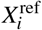 the position of MG atom *i* in the predicted and native structures. Both coordinate sets are evaluated after superposing the protein residues with an atom within 4 Å of the native MG. The resulting RMSD measures the MG pose relative to its binding pocket.

PPI predictions with DockQ > 0.23 are classified as acceptable in the CAPRI assessment^35^, with 0.49 and 0.80 marking the medium- and high-quality bands. MG accuracy was reported as the fractions with LDDT-PLI above 0.5 and 0.8 and ligand RMSD below 2, 5 and 10 Å. LDDT-PLI above 0.5 and ligand RMSD below 2 Å were used to assess recovery of the binding mode^22^. DockQ was computed with version 2.1.3 of the DockQ package^46^. LDDT-PLI and pocket-aligned ligand RMSD were computed using OpenStructure’s validation toolkit^47^. For the docking methods, predicted protein traces were completed by transferring native side chains rigidly onto the predicted C_*α*_ positions before scoring.

We used the original MGBench homology assignments, based on MMseqs2 searches against protein sequences from PDB structures released before the AF3 cutoff^22^. For complexes with two partners of at least 16 residues, high sequence homology requires both partners to exceed 40% sequence identity to two chains within the same reference complex. When one partner has fewer than 16 residues, inclusion in the subset with low sequence homology requires the longer partner to have less than 40% identity to every protein chain in this reference set. These criteria yield 25 complexes with low sequence homology and 63 with high sequence homology.

We generated *N* independent samples per complex across both choices of fixed protein and evaluated three prediction-selection rules. The top-1 pose is the highest-scoring candidate selected by the confidence model for use at inference. The blind pose is a single sample with the longer protein fixed. When sampling yields no structure, the next sampling round supplies the prediction. The best-of-*N* pose is the most accurate candidate according to the native structure. This oracle measure defines the maximum PPI prediction success rate attainable by ranking the same samples. The curves against *N* average over the whole set of samples and count a run that produces no structure as a failure.

For the cofolding comparison, we used versions of AF3, OpenFold3, ESMFold2, Protenix-v1 and Protenix-v2 with a PDB training cutoff of 30 September 2021, matching the cutoff used to construct Ternary-PDB. For MGBench, we compared DiffTernary with AF3, OpenFold3, ESMFold2, Protenix-v1 and Protenix-v2^12,17,19–21^, using 25 predictions per complex for each method. Cofolding methods received protein sequences and the MG chemical structure, while DiffTernary received the bound protein conformations and MG chemical structure. All predictions were scored against the same experimental structures using the same evaluation procedure, and best-of-25 performance was determined by the highest DockQ for each complex. We also generated DeepTernary predictions using the released MGD model weights^9^. Each 25-sample docking set comprised 13 samples with the longer protein fixed and 12 with the shorter protein fixed. Single predictions used the first successful sample with the longer protein fixed. Seven MGBench PDB entries appeared in the released DeepTernary model’s training list, including five with the same protein-chain and MG identifiers. All 88 MGBench complexes were retained in the comparison. For the TernaryDB MGD test set, we compared DiffTernary, DeepTernary^9^, EquiDock^37^ and DiffDock-PP^28^ on the same 94 complexes. DiffTernary was trained on the TernaryDB MGD training split, and DeepTernary used its released MGD model weights. EquiDock^37^ and DiffDock-PP^28^ used their released checkpoints, trained on the DIPS database of interacting protein structures^48^, and their published inference settings. These protein–protein docking methods received the two proteins alone, with the longer protein fixed, from the same input files used for DeepTernary and DiffTernary. We evaluated single blind predictions to compare baseline docking accuracy and sample sets of up to 40 poses per complex to assess oracle performance. The DiffTernary confidence model was evaluated by selecting its highest-scoring pose from each sample set.

Runtime was measured over the full test split on the hardware listed in Table S3. Differences between methods were assessed using two-sided Wilcoxon signed-rank tests paired by complex, with Holm correction within each comparison family.

### E2Former score backbone

The E2Former variant replaces the four interaction layers with equivariant attention blocks. Wigner 6 *j* recoupling computes geometric tensor products at nodes for reuse across their edges^30^. We adapted these blocks to the ternary graph by keeping even and odd features separate and computing attention over each node’s full incoming neighborhood. Relation embeddings distinguish interactions within proteins, within the MG, between proteins and the MG, and between the two proteins.

Each node carries scalar and vector features with 32 channels for each angular degree and parity. Writing *p* +1, 1 for parity, the representation and its transformation under rotation are

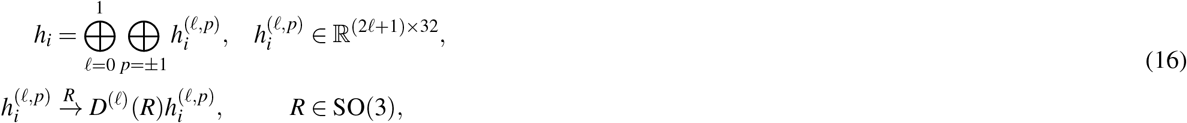

where *D*^(ℓ)^ is the degree-ℓ rotation matrix. Coupling a feature of parity *p*_in_ with a degree-*k* spherical harmonic gives *p*_out_ = *p*_in_( −1)^*k*^. Four attention heads combine messages over each node’s incoming edges. Attention weights depend on node features, distance, diffusion time and relation type. Even and odd features use separate channel maps, and attention scores are formed by contracting features of matching parity. Each block combines attention, an equivariant feed-forward network, normalization and residual connections. In the feed-forward branch, odd features are modulated by even scalar gates through bias-free linear maps. Normalization preserves the sign change of odd scalar features under inversion.

For a first-order angular coupling, the message to node *i* is

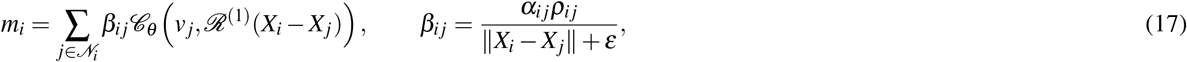

where *α*_*i j*_ is the attention weight normalized over incoming edges, *ρ*_*i j*_ is a learned radial modulation, *v* _*j*_ is the value feature, and *ε* = 10^−8^. The operator *C*_*θ*_ couples the value feature with the first-degree solid spherical harmonic *ℛ* ^(1)^. Angular-degree, parity and attention-head indices are omitted for clarity. Linearity of the solid harmonic gives

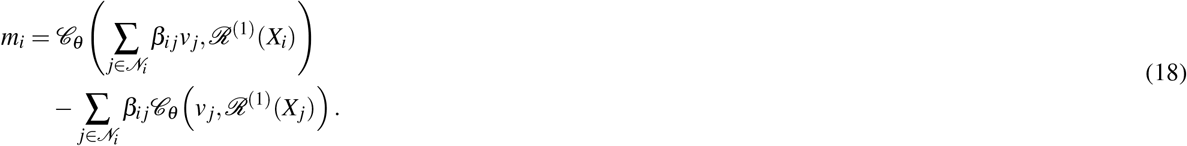

The sender coupling is computed once per node and reused across outgoing edges. The receiver coupling is computed after the weighted values have been summed. Attention and aggregation operate directly on the full edge list. After four blocks, the complete scalar and vector features are passed to the center graph for translation and rotation prediction (eq. (5)), followed by the magnitude mapping described above (eq. (6)). For torsion prediction, features are projected onto the original representation basis and passed through the bond-centered head (eq. (7)), with layer normalization before the final MLP.

### E2Former training and efficiency evaluation

The final E2Former model followed the training procedure described above, with the batch size increased to 16 per GPU and the learning rate to 0.002. Training throughput was evaluated using FP32 on eight H800 GPUs. The Tensor Product timings were taken from the main training log over 51 complete epochs, using the interval from the first to the last logged training batch. E2Former was measured in a separate timing run with a learning rate of 0.001 over two training epochs. Each epoch contained 42,328 training samples. Each training sample was a noisy ternary-complex graph with a specified choice of fixed protein. Throughput was calculated from the median epoch duration, with per-GPU batch sizes of 1 for Tensor Product and 16 for E2Former. Training-set evaluation passes were excluded from the timing summary.

The timed sampling comparison used the Tensor Product epoch-42 EMA checkpoint and the E2Former epoch-45 EMA checkpoint, with random seed 42 and *T* = 4. Both models used 20 time points and 19 updates, with no added noise in the final update. Every test complex was sampled under both choices of fixed protein, with 20 candidates per assignment. Blind prediction used the first successfully generated candidate with the longer protein fixed. Oracle prediction selected the highest-DockQ candidate from the combined pool of 40 candidate slots. All 94 complexes were retained. For the time-dependent curves, each completed sampling round added one candidate under each choice of fixed protein. Candidate quality was calculated over the accumulated pool of 2 to 40 candidates per complex in generation order.

Sampling used one A800 GPU, with FP32 and batch size 1 for Tensor Product and BF16 with batch and chunk sizes of 16 for E2Former. Timing used CUDA synchronization after warmup and included network evaluation, graph construction, pose updates and retries. Loading, structure scoring and output were excluded. Total sampling time was accumulated across the 20 sampling rounds. The average time per candidate was calculated over all 3,760 candidate slots (Table S1).

## Acknowledgements

We thank Peijie Zhou and Xudong Zhang from Peking University for helpful discussions and suggestions on DiffTernary model training.

## Author Contributions

Y.L. and J.Z. designed the research. Y.L. developed most of the code, conducted most of the experiments, and curated Ternary-PDB. L. Lin developed the E2Former variant. Y.L., L. Lin, J.Z., L. Lai and J.P. discussed the results. Y.L. and L. Lin wrote the manuscript. J.P., L. Lai and J.Z. supervised the project. J.P., L. Lai and J.Z. revised the manuscript. All authors read and approved the final manuscript.

## Supplementary Figures

**Figure S1.**
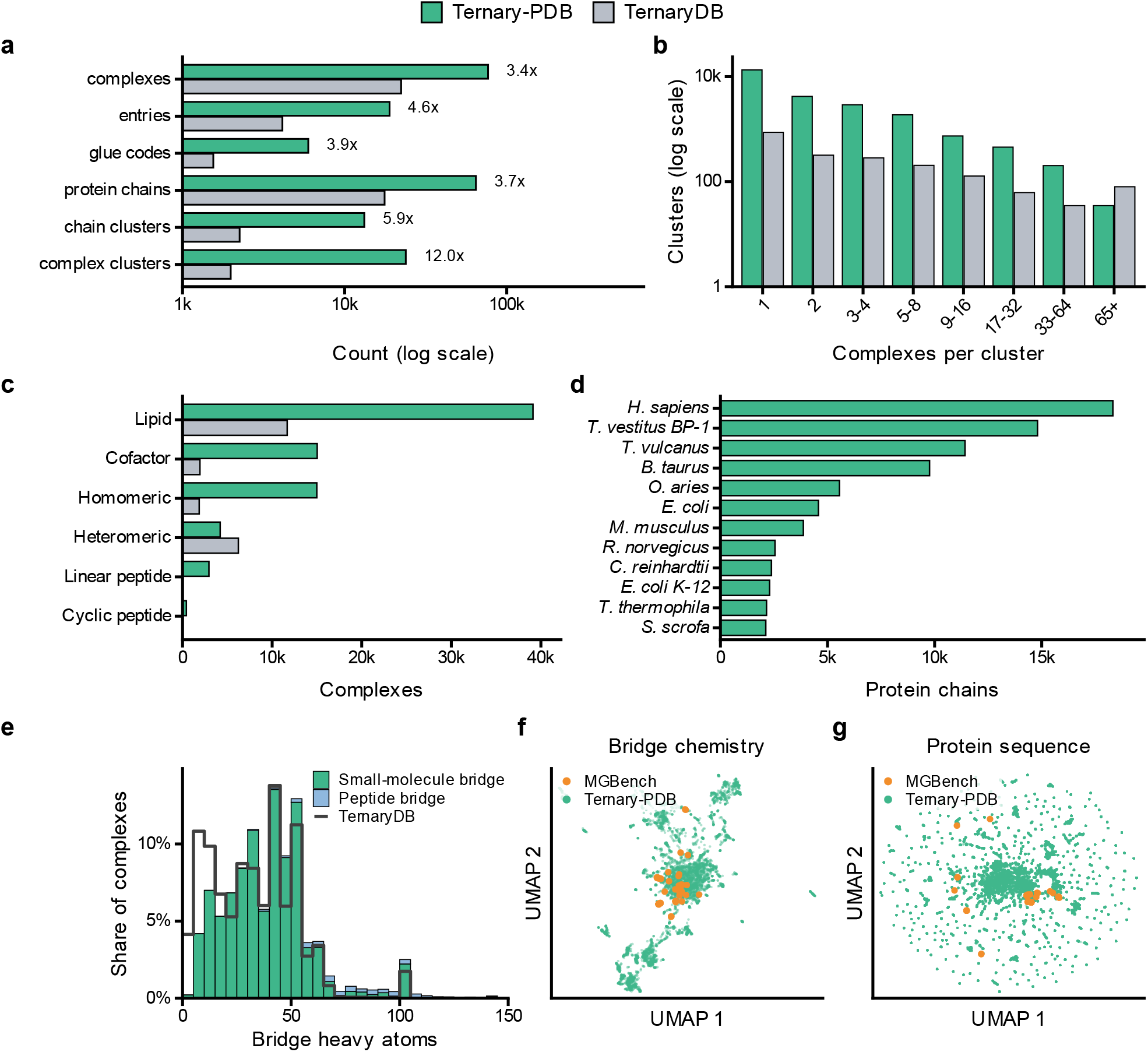
Dataset statistics and diversity. **a**, Dataset sizes and their ratios. **b**, Complex-cluster size distributions at 40% sequence identity for Ternary-PDB and 50% for TernaryDB. **c**, Bridge classes. **d**, Protein-chain occurrences from the twelve most represented organisms. **e**, Bridge heavy-atom counts in bins of five, separating small-molecule and peptide bridges. **f**, UMAP of 1024-bit Morgan fingerprints for 7,316 Ternary-PDB and 68 MGBench molecules, using radius 2 and Jaccard distance. **g**, UMAP of paired protein ESM2 embeddings using cosine distance. Ternary-PDB is green, TernaryDB gray and MGBench orange.

**Figure S2.**
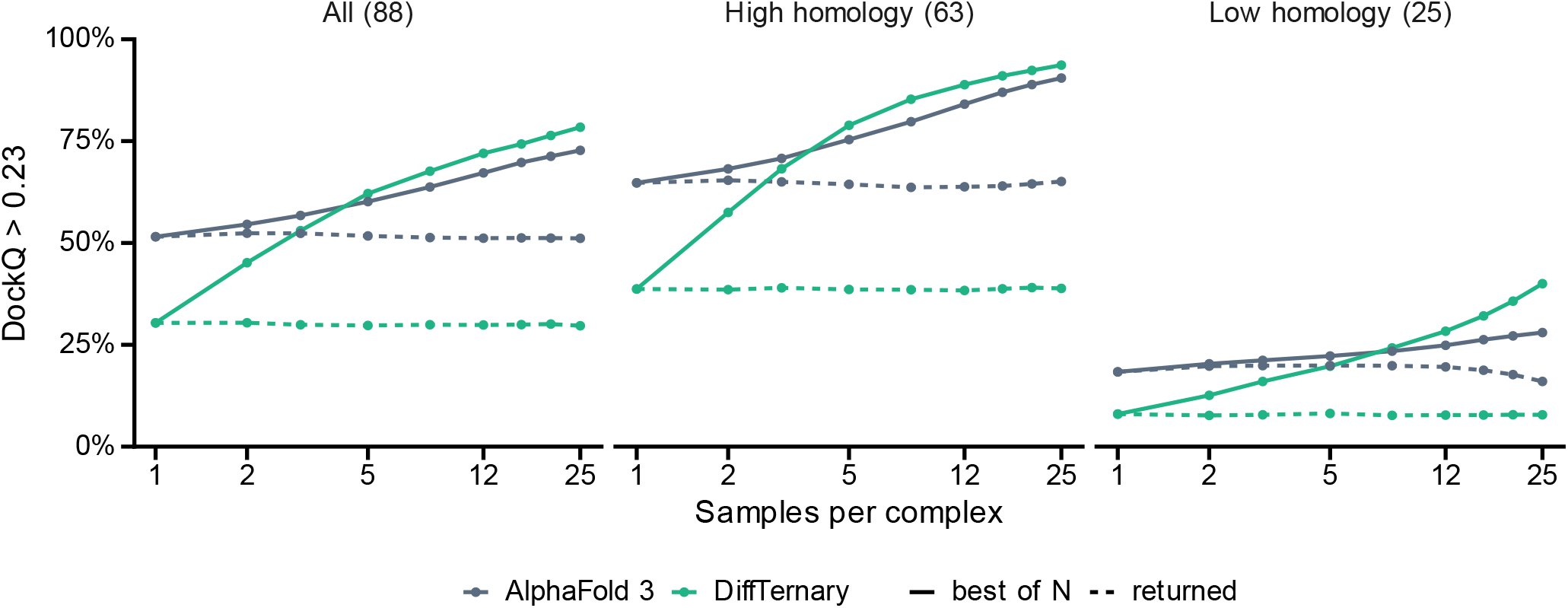
Sampling and pose selection on MGBench. Solid lines show best-of-*N* PPI prediction success rates. Broken lines show AF3 confidence selection or a single DiffTernary prediction. Curves average 400 random subsets of 25 attempts with seed 0. Missing predictions count as failures. DiffTernary uses ep116, with single-sample points averaged over both choices of fixed protein. The main blind comparison uses the first successful prediction with the longer protein fixed.

**Figure S3.**
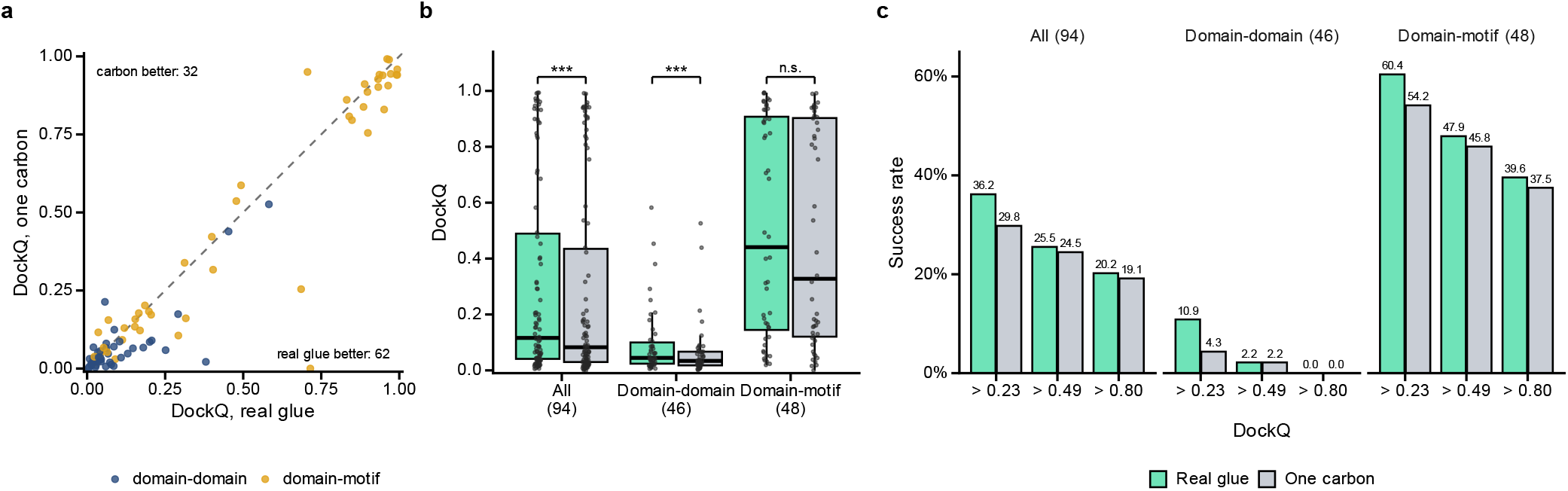
Docking with the MG or a single carbon atom. Each of 94 complexes was predicted with its MG or a carbon atom at the MG centroid. **a**, Paired DockQ values, colored by interface class. **b**, DockQ distributions overall and by class. **c**, PPI prediction success rates at DockQ thresholds of 0.23, 0.49 and 0.80. Two-sided paired Wilcoxon tests compare the conditions. Symbols denote \*\*\**P* < 0.001 and n.s. for a non-significant difference.

**Figure S4.**
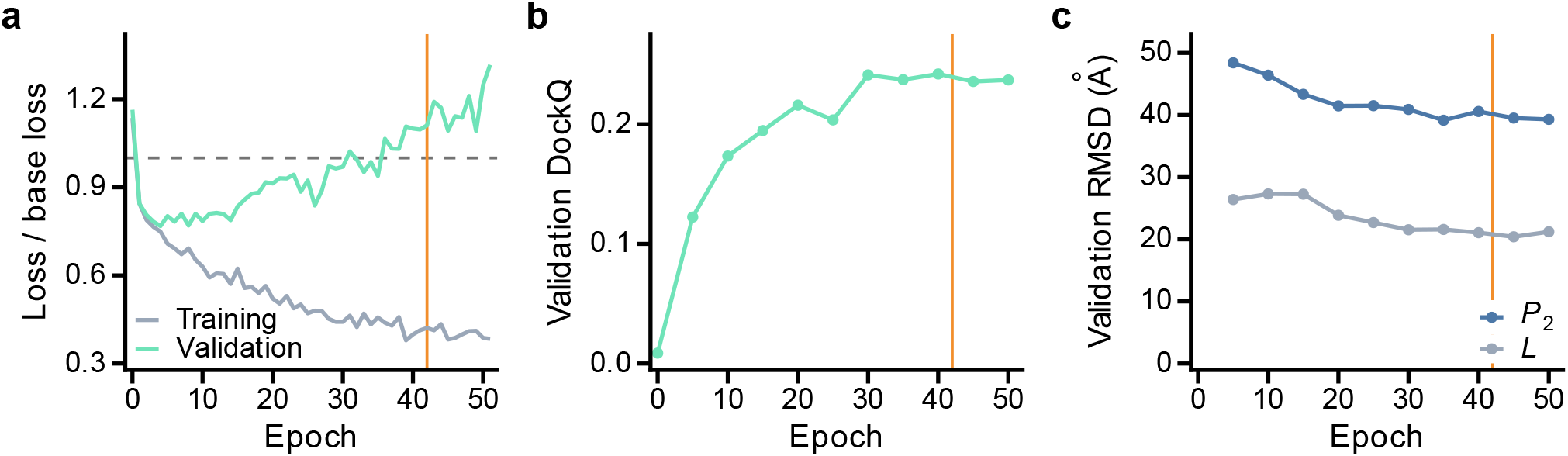
Score-model training on TernaryDB. **a**, Score matching loss divided by the zero-prediction baseline, marked by the dashed line. **b**, Mean validation DockQ, evaluated every five epochs on 137 complexes under both choices of fixed protein. **c**, Validation RMSD of the movable protein and MG. Vertical lines mark the epoch-42 weights used for evaluation.

**Figure S5.**
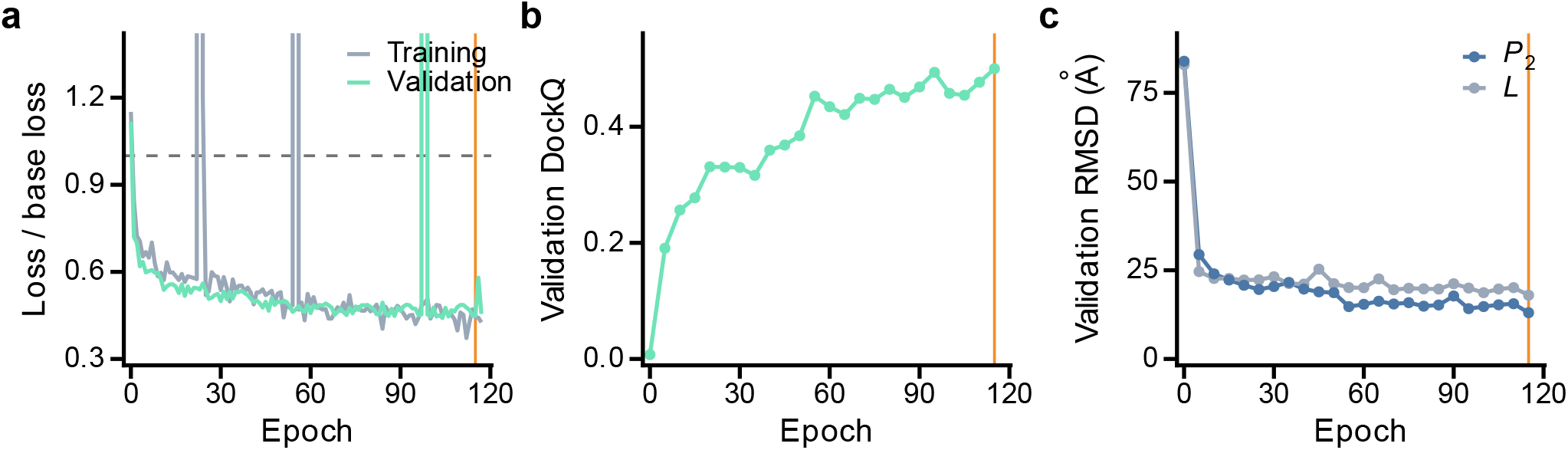
Score-model training on Ternary-PDB. Each epoch contains 42,328 training samples. **a**, Loss relative to the zero-prediction baseline (dashed line). Loss ratios outside the axis range and one non-finite validation value are not shown. **b**, Mean validation DockQ, evaluated every five epochs on 150 complexes under both choices of fixed protein. **c**, Validation RMSD of the movable protein and MG. Vertical lines mark epoch index 115, corresponding to ep116. Loss measurements extend through epoch 117 and docking evaluations through epoch 115.

**Figure S6.**
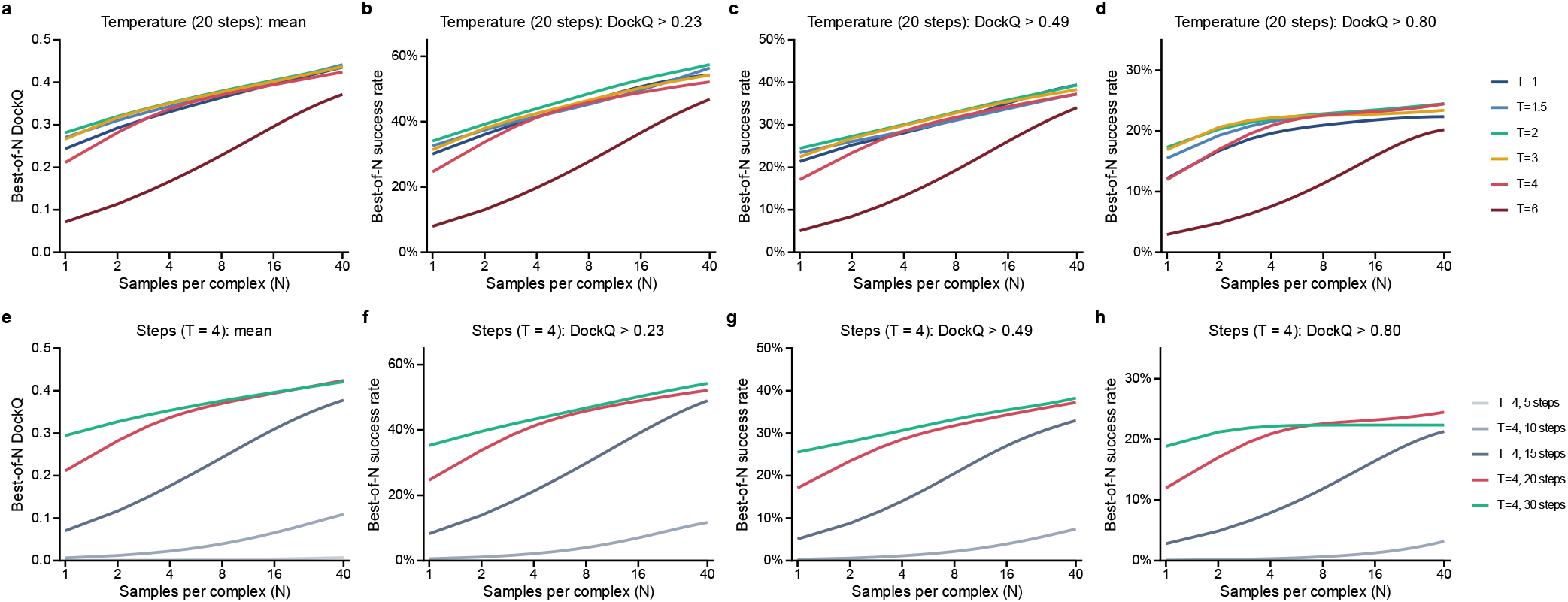
Sampling temperature and denoising steps. Best-of-*N* PPI prediction success rates on 94 TernaryDB complexes. **a–d**, Six temperatures with 20 denoising steps. **e–h**, Five step counts at *T* = 4.

## Supplementary Tables

**Table S1.** Tensor Product and E2Former efficiency. Training times summarize 51 Tensor Product epochs and two E2Former timing epochs, each containing 42,328 samples. Sampling covers 94 complexes with 20 samples per choice of fixed protein. Blind selection uses the first successful sample with the longer protein fixed and oracle selection the best of 40. Sampling times include network evaluation, graph and pose updates and retries after warmup, excluding loading, scoring and output.

| Metric | Tensor Product | E2Former |
|---|---|---|
| *Training timing* |  |  |
| Hardware | 8 × H800 | 8 × H800 |
| Precision | FP32 | FP32 |
| Batch size per GPU | 1 | 16 |
| Peak GPU memory | 74.78 GiB | 51.22 GiB |
| Learning rate in timing experiment | 0.001 | 0.001 |
| Measured training epochs | 51 | 2 |
| Median epoch duration | 3,310.00 s | 479.43 s |
| Training throughput | 12.79 samples/s | 88.29 samples/s |
| *Sampling timing* |  |  |
| Hardware | 1 × A800 | 1 × A800 |
| Checkpoint epoch | 42 | 45 |
| Precision | FP32 | BF16 |
| Batch size | 1 | 16 |
| Samples per complex | 40 | 40 |
| Total sampling time | 4,167.14 s | 952.71 s |
| Average sampling time per sample | 1.108 s | 0.253 s |
| Average sampling time per 40-sample set | 44.33 s | 10.14 s |
| *Blind prediction, 94 complexes* |  |  |
| Mean DockQ | 0.2577 | 0.2358 |
| Median DockQ | 0.0796 | 0.0513 |
| DockQ >0.23 | 30.9% (29/94) | 29.8% (28/94) |
| DockQ >0.49 | 21.3% (20/94) | 19.1% (18/94) |
| DockQ >0.80 | 14.9% (14/94) | 16.0% (15/94) |
| *Best of 40, 94 complexes* |  |  |
| Mean DockQ | 0.4237 | 0.4082 |
| Median DockQ | 0.2604 | 0.2313 |
| DockQ >0.23 | 56.4% (53/94) | 50.0% (47/94) |
| DockQ >0.49 | 37.2% (35/94) | 36.2% (34/94) |
| DockQ >0.80 | 22.3% (21/94) | 23.4% (22/94) |

**Table S2.** Tensor Product score and confidence model settings. Shared settings span both columns. Parameter counts exclude buffers.

|  | Score model | Confidence model |
|---|---|---|
| *Graph* |  |  |
| Protein node | One per residue at its C<sub>α</sub> atom |  |
| Protein node features | Amino-acid type and the 1280-dimensional ESM2-650M embedding |  |
| MG node | One per heavy atom |  |
| Protein edges | 30 Å, at most 24 neighbors |  |
| MG edges | Covalent bonds and a 5 Å radius graph |  |
| MG to protein edges | 3σ<sub>tr</sub>(*t*) + 20 Å, at *t* = 0 for the confidence model |  |
| Protein to protein edges | 3σ<sub>tr</sub>(*t*) + 40 Å, at *t* = 0 for the confidence model |  |
| Radial expansion | 32 Gaussians within a component, 32 across components |  |
| Time embedding | Sinusoidal, dimension 32, scale 10,000 |  |
| *Network* |  |  |
| Scalar channels *n<sub>s</sub>* | 32 |  |
| Vector channels *n<sub>v</sub>* | 6 |  |
| Spherical harmonics | ℓ ≤ 2, first order representations only |  |
| Embedding layers | 1 per component |  |
| Interaction layers | 4 |  |
| Edge groups per layer | 4 |  |
| Tensor-product weight network | Two layers, width 3*n<sub>s</sub>* |  |
| Batch normalization | Off |  |
| Dropout | 0.1 |  |
| Parameters | 4,420,602 | 4,116,625 |
| *Diffusion* |  |  |
| σ<sub>tr</sub> | 0.1 to 30 Å |  |
| σ<sub>rot</sub> | 0.03 to 1.55 rad |  |
| σ<sub>tor</sub> | 0.0314 to 3.14 rad |  |
| Interpolation in *t* | Exponential |  |
| Loss weights, MG | 0.167, 0.167, 0.166 |  |
| Loss weights, partner | 0.25, 0.25 |  |
| *Optimization* |  |  |
| Optimizer | Adam, learning rate 10<sup>-3</sup>, no weight decay |  |
| Scheduler | Plateau, patience 10 | Plateau, patience 30 |
| Gradient clipping | Norm 1 |  |
| Weight averaging | Exponential, rate 0.999 | None |
| Batch size per rank | 1 | 4 |
| Checkpoint selection | Validation DockQ | Validation interface RMSD of the top-ranked pose |
| *Inference* |  |  |
| Denoising schedule | 20 time points, 19 updates |  |
| Final step | Taken without added noise |  |
| Sampling temperature | *T* = 4, no churn |  |
| Samples per complex | 40 |  |

**Table S3.**
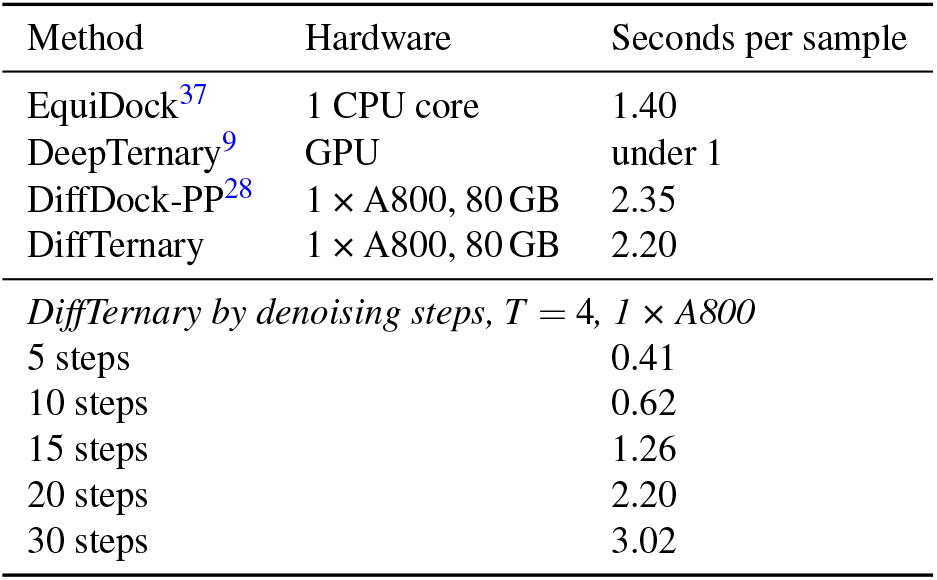
Average time per sample on TernaryDB. Full-run wall-clock time, including startup, loading, sampling and ranking where applicable, divided by the total samples across 94 complexes. DiffDock-PP and DiffTernary generated 40 samples per complex, and EquiDock and DeepTernary one. A800 runs processed one complex at a time. DeepTernary timing is from its authors. The lower block varies denoising steps at *T* = 4.

| Method | Hardware | Seconds per sample |
|---|---|---|
| EquiDock<sup>37</sup> | 1 CPU core | 1.40 |
| DeepTernary<sup>9</sup> | GPU | under 1 |
| DiffDock-PP<sup>28</sup> | 1 × A800, 80 GB | 2.35 |
| DiffTernary | 1 × A800, 80 GB | 2.20 |
| *DiffTernary by denoising steps, T* = 4*, 1* × *A800* |  |  |
| 5 steps |  | 0.41 |
| 10 steps |  | 0.62 |
| 15 steps |  | 1.26 |
| 20 steps |  | 2.20 |
| 30 steps |  | 3.02 |

## References

1. Schreiber, S. L. Molecular glues and bifunctional compounds: Therapeutic modalities based on induced proximity. Cell Chem. Biol. 31, 1050–1063, DOI: 10.1016/j.chembiol.2024.05.004 (2024).

2. Konstantinidou, M. & Arkin, M. R. Molecular glues for protein-protein interactions: progressing toward a new dream. Cell Chem. Biol. 31, 1064–1088, DOI: 10.1016/j.chembiol.2024.04.002 (2024).

3. Rui, H., Ashton, K. S., Min, J., Wang, C. & Potts, P. R. Protein–protein interfaces in molecular glue-induced ternary complexes: classification, characterization, and prediction. RSC Chem. Biol. 4, 192–215, DOI: 10.1039/d2cb00207h (2023).

4. Zhu, J., Lai, L. & Pei, J. Toward in silico design of protein–protein interaction stabilizers. ACS Cent. Sci. 9, 861–863, DOI: 10.1021/acscentsci.3c00545 (2023).

5. Zhu, J. et al. MGTbind: a comprehensive database of molecular glue ternary interactome. Nucleic Acids Res. 54, D1500–D1509, DOI: 10.1093/nar/gkaf1075 (2026).

6. Kozicka, Z. & Thomä, N. H. Haven’t got a glue: Protein surface variation for the design of molecular glue degraders. Cell Chem. Biol. 28, 1032–1047, DOI: 10.1016/j.chembiol.2021.04.009 (2021).

7. Mayor-Ruiz, C. et al. Rational discovery of molecular glue degraders via scalable chemical profiling. Nat. Chem. Biol. 16, 1199–1207, DOI: 10.1038/s41589-020-0594-x (2020).

8. Robinson, S. A., Co, J. A. & Banik, S. M. Molecular glues and induced proximity: An evolution of tools and discovery. Cell Chem. Biol. 31, 1089–1100, DOI: 10.1016/j.chembiol.2024.04.001 (2024).

9. Xue, F. et al. SE(3)-equivariant ternary complex prediction towards target protein degradation. Nat. Commun. 16, 5514, DOI: 10.1038/s41467-025-61272-5 (2025).

10. Lin, H. et al. Leveraging high-throughput proteomics and AI-based protein folding to accelerate VAV1 molecular glue discovery. Nat. Commun. 17, 10763, DOI: 10.1038/s41467-026-77657-z (2026).

11. Krishna, R. et al. Generalized biomolecular modeling and design with RoseTTAFold All-Atom. Science 384, eadl2528, DOI: 10.1126/science.adl2528 (2024).

12. Abramson, J. et al. Accurate structure prediction of biomolecular interactions with AlphaFold 3. Nature 630, 493–500, DOI: 10.1038/s41586-024-07487-w (2024).

13. Chai Discovery et al. Chai-1: decoding the molecular interactions of life. bioRxiv 2024.10.10.615955, DOI: 10.1101/2024.10.10.615955 (2024). Version 2.

14. Wohlwend, J. et al. Boltz-1: democratizing biomolecular interaction modeling. bioRxiv 2024.11.19.624167, DOI: 10.1101/2024.11.19.624167 (2024). Version 4, updated 6 May 2025.

15. Passaro, S., Corso, G., Wohlwend, J., Reveiz, M. et al. Boltz-2: Towards accurate and efficient binding affinity prediction. bioRxiv DOI: 10.1101/2025.06.14.659707 (2025).

16. ByteDance AML AI4Science Team et al. Protenix - advancing structure prediction through a comprehensive AlphaFold3 reproduction. bioRxiv 2025.01.08.631967, DOI: 10.1101/2025.01.08.631967 (2025).

17. The OpenFold3 Team. OpenFold3-preview, DOI: 10.5281/zenodo.19001000 (2025). Research preview, version 0.4.2, official repository citation.

18. Aureka AI OpenDDE project. Folding, reasoning, and scaling with Open-source Drug Discovery Engine, DOI: 10.48550/arXiv.2607.03787 (2026). 2607.03787.

19. Candido, S. et al. Language modeling materializes a world model of protein biology. bioRxiv DOI: 10.64898/2026.06.03.729735 (2026).

20. Protenix Team et al. Protenix-v1: Toward high-accuracy open-source biomolecular structure prediction. bioRxiv DOI: 10.64898/2026.02.05.703733 (2026).

21. Zhang, Y. et al. Protenix-v2: Broadening the reach of structure prediction and biomolecular design. bioRxiv DOI: 10.64898/2026.04.10.717613 (2026).

22. Liao, Y., Zhu, J., Xie, J., Lai, L. & Pei, J. Benchmarking cofolding methods for molecular glue ternary structure prediction. J. Chem. Inf. Model. 65, 11136–11148, DOI: 10.1021/acs.jcim.5c01860 (2025).

23. Zaidman, D., Prilusky, J. & London, N. PRosettaC: Rosetta based modeling of PROTAC mediated ternary complexes. J. Chem. Inf. Model. 60, 4894–4903, DOI: 10.1021/acs.jcim.0c00589 (2020).

24. Drummond, M. L., Henry, A., Li, H. & Williams, C. I. Improved accuracy for modeling PROTAC-mediated ternary complex formation and targeted protein degradation via new in silico methodologies. J. Chem. Inf. Model. 60, 5234–5254, DOI: 10.1021/acs.jcim.0c00897 (2020).

25. Weng, G., Li, D., Kang, Y. & Hou, T. Integrative modeling of PROTAC-mediated ternary complexes. J. Med. Chem. 64, 16271–16281, DOI: 10.1021/acs.jmedchem.1c01576 (2021).

26. Corso, G., Stärk, H., Jing, B., Barzilay, R. & Jaakkola, T. DiffDock: Diffusion steps, twists, and turns for molecular docking. In International Conference on Learning Representations (2023).

27. Zhu, J., Gu, Z., Pei, J. & Lai, L. DiffBindFR: an SE(3) equivariant network for flexible protein–ligand docking. Chem. Sci. 15, 7926–7942, DOI: 10.1039/D3SC06803J (2024).

28. Ketata, M. A. et al. DiffDock-PP: Rigid protein-protein docking with diffusion models. In ICLR Machine Learning for Drug Discovery Workshop (2023).

29. Wang, Y., Wang, F., Feng, L., Zhang, C. & Lai, L. DiffPepDock: Efficient protein–peptide docking and binder screening via SE(3)-equivariant diffusion. Protein Sci. 34, e70338, DOI: 10.1002/pro.70338 (2025).

30. Li, Y. et al. E2Former: An efficient and equivariant transformer with linear-scaling tensor products. In Advances in Neural Information Processing Systems, vol. 38, 26777–26814, DOI: 10.52202/085713-0797 (2025).

31. Berman, H. M. et al. The Protein Data Bank. Nucleic Acids Res. 28, 235–242, DOI: 10.1093/nar/28.1.235 (2000).

32. Steinegger, M. & Söding, J. MMseqs2 enables sensitive protein sequence searching for the analysis of massive data sets. Nat. Biotechnol. 35, 1026–1028, DOI: 10.1038/nbt.3988 (2017).

33. Landrum, G. & RDKit contributors. RDKit: open-source cheminformatics. https://www.rdkit.org, DOI: 10.5281/zenodo.15605628 (2025). Version 2025.03.3.

34. Basu, S. & Wallner, B. DockQ: a quality measure for protein-protein docking models. PLoS One 11, e0161879, DOI: 10.1371/journal.pone.0161879 (2016).

35. Méndez, R., Leplae, R., Lensink, M. F. & Wodak, S. J. Assessment of CAPRI predictions in rounds 3–5 shows progress in docking procedures. Proteins 60, 150–169, DOI: 10.1002/prot.20551 (2005).

36. Singhal, R. et al. A general framework for inference-time scaling and steering of diffusion models. In Proceedings of the 42nd International Conference on Machine Learning, vol. 267 of Proceedings of Machine Learning Research, 55810–55827 (PMLR, 2025).

37. Ganea, O.-E. et al. Independent SE(3)-equivariant models for end-to-end rigid protein docking. In International Conference on Learning Representations (2022).

38. Corso, G., Deng, A., Polizzi, N., Barzilay, R. & Jaakkola, T. Deep confident steps to new pockets: Strategies for docking generalization. In International Conference on Learning Representations (ICLR) (2024).

39. Vakser, I. A. Protein-protein docking: from interaction to interactome. Biophys. J. 107, 1785–1793, DOI: 10.1016/j.bpj.2014.08.033 (2014).

40. Nikolayev, D. I. & Savyolova, T. I. Normal distribution on the rotation group SO(3). Textures Microstruct. 29, 201–233, DOI: 10.1155/TSM.29.201 (1997).

41. Leach, A., Schmon, S. M., Degiacomi, M. T. & Willcocks, C. G. Denoising diffusion probabilistic models on SO(3) for rotational alignment. In ICLR Workshop on Geometrical and Topological Representation Learning (2022).

42. Jing, B., Corso, G., Chang, J., Barzilay, R. & Jaakkola, T. Torsional diffusion for molecular conformer generation. In Advances in Neural Information Processing Systems, vol. 35, 24240–24253, DOI: 10.52202/068431-1760 (2022).

43. Geiger, M. & Smidt, T. e3nn: Euclidean neural networks. arXiv preprint 2207.09453 DOI: 10.48550/arXiv.2207.09453 (2022).

44. Lin, Z. et al. Evolutionary-scale prediction of atomic-level protein structure with a language model. Science 379, 1123–1130, DOI: 10.1126/science.ade2574 (2023).

45. Robin, X. et al. Assessment of protein–ligand complexes in CASP15. Proteins 91, 1811–1821, DOI: 10.1002/prot.26601 (2023).

46. Mirabello, C. & Wallner, B. DockQ v2: improved automatic quality measure for protein multimers, nucleic acids, and small molecules. Bioinformatics 40, btae586, DOI: 10.1093/bioinformatics/btae586 (2024).

47. Biasini, M. et al. OpenStructure: an integrated software framework for computational structural biology. Acta Crystallogr. D Biol. Crystallogr. 69, 701–709, DOI: 10.1107/S0907444913007051 (2013).

48. Townshend, R. J. L., Bedi, R., Suriana, P. A. & Dror, R. O. End-to-end learning on 3D protein structure for interface prediction. In Advances in Neural Information Processing Systems, vol. 32 (2019).

